# Phage-mediated resolution of genetic conflict alters the evolutionary trajectory of *Pseudomonas aeruginosa* lysogens

**DOI:** 10.1101/2023.10.24.563318

**Authors:** Laura C. Suttenfield, Zoi Rapti, Jayadevi H. Chandrashekhar, Amelia C. Steinlein, Juan Cristobal Vera, Ted Kim, Rachel J. Whitaker

## Abstract

The opportunistic human pathogen *Pseudomonas aeruginosa* is naturally infected by a large class of temperate, transposable, Mu-like phages. We examined the genotypic and phenotypic diversity of *P. aeruginosa* PA14 populations as they resolve CRISPR autoimmunity, mediated by an imperfect CRISPR match to the Mu-like DMS3 prophage, and show that lysogen evolution is profoundly impacted by CRISPR autoimmunity and phage transposition around the chromosome. After 12 days of evolution, we measured a decrease in spontaneous induction in both exponential and stationary phase growth. Co-existing variation in spontaneous induction rates in exponential phase corresponded to a difference in the type of CRISPR self-targeting resolution, mediated either by host mutation or phage transposition. Multiple mutational modes to resolve genetic conflict between host and phage resulted in coexistence in evolved populations of single lysogens that maintained CRISPR immunity to other phages and polylysogens that have lost immunity completely. This work highlights a new dimension of the role of lysogenic phages in the evolution of their hosts.

**Importance:** The chronic opportunistic multi-drug resistant pathogen *Pseudomonas aeruginosa* is persistently infected by temperate phages. We assess the contribution of temperate phage infection to the evolution of the clinically relevant strain UCBPP-PA14. We found that a low level of CRISPR-mediated self-targeting resulted in polylysogeny evolution and large genome rearrangements in lysogens; we also found extensive diversification in CRISPR spacers and *cas* genes. These genomic modifications resulted in decreased spontaneous induction in both exponential and stationary phase growth, increasing lysogen fitness. This work shows the importance of considering latent phage infection in characterizing the evolution of bacterial populations.

## Introduction

Cystic fibrosis (CF) is a genetic disorder which makes patients vulnerable to respiratory infections by commensal and environmental bacterial pathogens like *Pseudomonas aeruginosa*. *Pseudomonas aeruginosa* is a common human pathogen whose increase in multi-drug antibiotic resistance has made it the focus of targeted phage therapy (Kortright et al., 2019). In chronic *Pseudomonas* infections of CF patients, it is common to find that all isolates track their origin to a single ancestral genotype (Feliziani et al., 2014; Folkesson et al., 2012; Jorth et al., 2015; Markussen et al., 2014; Schick & Kassen, 2018; Tai et al., 2017; Workentine et al., 2013). Diversity of *P. aeruginosa* strains diverge from this common ancestor through *de novo* mutations generated by mutation and hypermutation genotypes (Vanderwoude et al., 2023), recombination (Darch et al., 2015; Vanderwoude et al., 2023) and large deletions (Cramer et al., 2011; Jeukens et al., 2014; Rau et al., 2012).

*Pseudomonas aeruginosa* is commonly infected by latent phages (bacteriophage) when colonizing CF patients (Budzik et al., 2004; Burgener et al., 2019; Folkesson et al., 2012; Ojeniyi et al., 1991; Rossi et al., 2021; Tariq et al., 2019; Vanderwoude et al., 2023; Winstanley et al., 2016). Temperate and chronic phages act as both sources of genetic novelty (Brüssow et al., 2004) and as potential assassins that can be induced to kill their hosts (James et al., 2015; Willner et al., 2012). In the lung environment, the presence of antibiotics and reactive oxygen species (Kettle et al., 2014; Malhotra et al., 2019; McGrath et al., 1999) may act as inducing agents for phage (Bondy-Denomy et al., 2016; Fothergill et al., 2011; James et al., 2015; Nanda et al., 2015; Rolain et al., 2009). Bacterial lysis through phage induction is hypothesized to help control bacterial growth in the lung (James et al., 2015) and may be used in synergy with antibiotics (Al-Anany et al., 2021; Clifton et al., 2019). Additionally, lysogeny has been shown to co-occur with host genome rearrangements in chronically infecting *Staphylococcus aureus* (Goerke et al., 2004, 2006; Golubchik et al., 2013; Guérillot et al., 2019), and *Streptococcus pyogenes* (Nakagawa et al., 2003). However, it remains unclear if and how lysogeny alters the evolution of the host genome.

Mu-like transposable phages are a diverse family of phages which infect an equally diverse range of bacteria (Zhang et al., 2023). Upon infection of the host, Mu-like phages integrate into the host chromosome through a conservative (cut-and-paste) transposition step that occurs with low sequence preference (Chaconas & Harshey, 2007; Vergnaud et al., 2018; Walker et al., 2020). Lytic replication occurs via replicative (copy-and-paste) transposition around the genome (Walker & Harshey, 2020), which occurs approximately 100 times and terminates in headful packaging into the virion. In lysogeny, which is established in approximately 10% of infections, a low-specificity insertion can increase the variation available to natural selection in *P. aeruginosa* populations through knockout mutations (Davies et al., 2016; O’Brien et al., 2019; Rollie et al., 2020). These *P. aeruginosa* lysogens have previously been found to have high viral titers in culture (Bondy-Denomy et al., 2016; James et al., 2012). Due to the nature of the chemistry that governs the transposition reaction, these insertions may also cause structural rearrangements such as deletions (Toussaint & Rice, 2017).

CRISPR-Cas (<u>c</u>lustered regularly interspaced <u>s</u>hort <u>p</u>alindromic repeats and <u>C</u>RISPR-<u>as</u>sociated genes) is a bacterial and archaeal adaptive immune system which incorporates foreign DNA fragments into an array as a spacer, and subsequently targets any piece of invading DNA (the protospacer) which is complementary to the spacer (Barrangou et al., 2007; Vorontsova et al., 2015). The Mu-like temperate phage are the most commonly targeted phages by CRISPR-Cas in *Pseudomonas aeruginosa* (England et al., 2018). Phage DMS3, a member of this group, was recovered from a *P. aeruginosa* CF isolate and infects the type strain UCBPP-PA14 (Budzik et al., 2004). DMS3 inhibits quorum sensing and pilus formation in PA14 lysogens (Shah et al., 2021). PA14 contains a Type 1-F CRISPR system with a partial spacer match to DMS3 (Zegans et al., 2009). This spacer has 5 mismatches to the phage protospacer, which is not sufficient to mediate immunity to the phage but leads to genetic conflict in DMS3 lysogens. This degenerate protospacer-spacer mismatch between DMS3 and PA14 targets the PA14 chromosome, causing enough DNA damage to stimulate the SOS response, which leads to the expression of pyocin genes, cell death, and limitation of biofilm formation (Heussler et al., 2015). Lysogens arising from PA14 cultures infected with free DMS3 virions evolved to have a lower spontaneous induction in stationary phase and lost their CRISPR systems over a seven-day evolutionary period. This was suggested to resolve genetic conflict caused by CRISPR self-targeting (immunopathology), a phenomenon which is predicted to be common in bacteria with Type 1 CRISPR systems and temperate phages (Rollie et al., 2020). However, how lysogeny alone impacts the bacterial genome during evolution in the absence of new phage infection has not yet been explored.

Here we directly assess the contribution of CRISPR-mediated genetic conflict between host and temperate phage to the evolution of *Pseudomonas* by analyzing evolved lysogen populations. We show that selection to resolve genetic conflict alters the evolutionary landscape of lysogen populations. Experimental work combined with genomic analysis demonstrates that transposable phages are a major source of variation beyond mutation that impacts the evolutionary direction of *P. aeruginosa* lysogens.

## Methods

### Experimental evolution

To establish the contribution of phages to host genome evolution, we evolved the uninfected laboratory strain UCBPP-PA14 (Rahme et al., 1995) and the established lysogen Lys2 (Zegans et al., 2009) for 12 days by serial transfer. Our strains are listed in Table S1. Lys2 is derived from PA14 and contains DMS3, which mediated a ∼20 kb host deletion from 806,169 to 826,108 bp on our PA14 reference chromosome, and a single A<G nonsynonymous point mutation at position 4,755,306 in a hypothetical protein (deleted genes are listed in Table S2). For each strain we randomly selected three colonies and grew up independent overnight cultures in LB (10 g tryptone; 5 g yeast extract; 5 g sodium chloride per liter of water). We subcultured the cells and normalized to an OD_600_ of 0.2 in 25 mL microcosms in three parallel 250 mL flasks. We serially passaged triplicate cultures with daily 1:100 transfers for 12 days, with shaking and at 37°C, for approximately 72 bacterial generations. At the end of the experiment, we colony-purified six colonies isolated from each replicate, and six colonies from the Lys2 ancestor stock for further analysis. Six isolates from each uninfected population were also sequenced and the data is presented in Supplementary Figure 1.

### One-step growth curves

In order to calculate the burst size of DMS3, we performed one-step growth curves (Ellis & Delbrück, 1939). Overnight, stationary phase PA14 cultures were diluted 1:100 and grown in LB until they reached an OD of 0.5. Phage was added to these cultures at a 1:10 volume ratio, for a final MOI of 0.01, mixed well, and incubated at 37°C for 5 minutes. In order to calculate adsorption, a fraction of the sample volume was immediately spun down for 5 mins at 30,000 rpm, and the supernatant was stored with 10% v/v chloroform to quantify the remaining free phage. To begin the one-step growth curve, the remainder of the samples were added to pre-warmed medium at a 1:100 ratio to stop adsorption, and grown on a roller drum at 37°C for the next 2.5 hours. Samples were taken every 10 minutes and mixed with a 10% v/v of chloroform for later quantification of PFUs.

We calculated burst size with the following equation: the total phage produced (the difference between the average PFUs before and after the burst) was divided by number of cells that were infected (estimated by the number of adsorbed phages multiplied by the estimated number of cells which proceeded through lysis). Our measurements of an 8% lysogenization frequency, estimated by spot-on-lawn assay, corresponded to measures in (Cady & O’Toole, 2011) (data not shown).

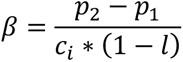

Here, *β* is burst size; *p_2_* and *p_1_* are the second and first plateaus, respectively; *c_i_* is the number of infected cells; *l* is the lysogenization frequency. The burst size of DMS3 is 41.8 +/− 8.4 phages per lysed cell (Fig S2). All PFUs were enumerated by spotting the phage-containing fraction on 0.5% double agar overlay plates.

### Spontaneous induction measurements

Because multiple inputs could contribute to a higher raw PFU value in stationary phase, and because stationary phase itself could be an inducing condition for some viruses (including phage Mu) (Ranquet et al., 2005), we chose to measure spontaneous induction in exponential phase, which necessitated the normalization of PFU values with the CFU values. This was also a desirable method for separating whether increased growth rate was responsible for increased PFU values. Due to these factors, we chose to measure spontaneous induction separately in both exponential and stationary phase. Our metric corresponds to other methods used to estimate these rates (Zong et al., 2010).

Growth curves were started from overnight cultures of replicate purified isolates. We washed cultures three times by resuspension in fresh LB media, normalized the OD to 0.2, and diluted them 1000-fold. Time points were taken at 0 and 2-7 hours to capture exponential phase growth, and plated for both CFUs and PFUs. Samples were incubated at 37°C on a roller drum. We measured spontaneous induction (*q*) by taking the difference in the number of viral particles released by cells growing in exponential phase, and normalized by the estimated burst size, the average growth of the culture, and the total time the culture grew.

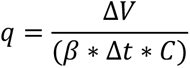

Here, *q* is spontaneous induction, and has units of burst cells per time; Δ*V* is the total increase in virion particles; *β* is the burst size; Δ*t* is the change in time; *C* is the average amount of cells in the culture. To account for exponential growth, all calculations were based on the linear regression of the log_10_ transformed Δ*V* and *C* values.

We also calculated spontaneous induction based on calculations in (Zong et al., 2010). This paper approximated spontaneous induction at each time point with this formula: 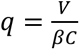; here, *V* is the number of virion particles at that time; *β* is the burst size; and *C* is the average amount of CFUs at that time. These methods produced qualitatively similar results (Fig S3).

To measure induction in stationary phase, we began growth curves in the same way as the exponential phase measurements. Time points were taken at 0, 10, 12, 15, and 18 hours, and plated for both CFUs and PFUs. To calculate the rate of spontaneous induction, we used the same formula as above, except with un-regressed logged values.

### Genome sequencing

In order to create a viral reference, DMS3 DNA was extracted from filtered Lys2 overnight supernatant using the Phage DNA Isolation Kit (Norgen Biotek Corp, Cat #: 46800), following the manufacturer’s instructions. Libraries were prepared with a Biomek 4000 liquid handler (Beckman-Coulter). We quantitated libraries with a Qubit fluorometer (Life Technologies Corporation, REF #: Q32866). Libraries were pooled and submitted for 2×250 paired-end sequencing by the Roy J. Carver Biotechnology Center at the University of Illinois Urbana-Champaign with an Illumina NovaSeq 6000. We received ∼3.8 million reads with about 100X coverage.

We inoculated evolved isolates and ancestral controls in 2 mL deep well plates in LB and grew them overnight. We extracted gDNA with the Beckman-Coulter gDNA extraction kit as above using the Nextera Flex Library Preparation Kit (Illumina). We quantitated libraries with a Qubit fluorometer (Life Technologies Corporation, REF #: Q32866). Libraries were pooled and submitted for 2165 cl:390250 paired-end sequencing by the Roy J. Carver Biotechnology Center at the University of Illinois Urbana-Champaign with an Illumina NovaSeq 6000. We received an average of about 5 million reads per genome. All raw reads are available on the NCBI database under BioProject number PRJNA1021667.

### Genome analysis

We ran a custom QC pipeline on our raw FASTQ reads, available on Github (http://www.github.com/igoh-illinois). Briefly, the Illumina adaptor sequences were trimmed using TrimmomaticPE v0. Read quality was checked with FastQC v0.11.9 (options: --noextract -k 5 -f fastq). Reads were aligned using BWA-MEM (Li, 2013) with default options to a 2-contig reference genome containing both our reference PA14 sequence and our reference DMS3 sequence. To identify chromosomal mutations, we ran Breseq (Deatherage & Barrick, 2014) on the trimmed and quality-controlled reads. SAM files were checked manually in both IGV 2.12.3 (J. T. Robinson et al., 2017) and Tablet (Milne et al., 2013).

To identify insertion sites of transposable phage, we ran a second pipeline, available on Github (http://www.github.com/igoh-illinois). The pipeline identifies insertion sites at nucleotide resolution by identifying reads that map to both the host and viral chromosome (“split” reads). It further identifies insertion sites by finding reads that have been split on either side of a 5-bp window, creating a small overlapping region when mapped back to the host genome. This is the result of a 5-bp duplication, which is characteristic of Mu and Mu-like phage transposition (Allet, 1979; Morgan et al., 2002).

While manually verifying the phage insertion sites in IGV, we found putative large duplicated regions of the host chromosome. To verify these duplicated regions we used the depth command in Samtools (Danecek et al., 2021) to find the number of reads that covered each position in the genome, and graphed this using RStudio (R version 4.3.1) (R Core Team, 2023). In order to assess the protein content of deleted and duplicated regions, FASTA files of sequence the reference PA14 genome was given to the eggNOG-mapper-v2 pipeline (settings: genomic data; default options) (Cantalapiedra et al., 2021).

We used the CRISPR Comparison Tool Kit (CCTK v1.0.0) to identify and compare CRISPR arrays (settings: crisprdiff; default options) (Collins & Whitaker, 2022).

### Mitomycin C induction experiments

Single colonies of the strains of interest were inoculated in LB, grown overnight at 37°C on a roller drum, and subcultured until they reached an OD of 0.5. Cultures were normalized, split, and incubated with or without 0.5 μg/μL mitomycin C (MMC) for 3.5 hours, after which CFUs and PFUs were enumerated.

### Model information

We use a compartmental model based on a system of ordinary differential equations. There are six lysogen compartments each representing a lysogen characterized by a distinct rate of spontaneous induction, but otherwise being identical. Lysogens are induced at their associated rates and transition into the lytic state which is followed by phage production and bursting. The model is along the lines of the ones presented and analyzed in our previous works (Clifton et al., 2019, 2021; Landa et al., 2021). All lysogens are assumed to grow at the same rate, and the total bacterial population grows logistically. The model equations read:

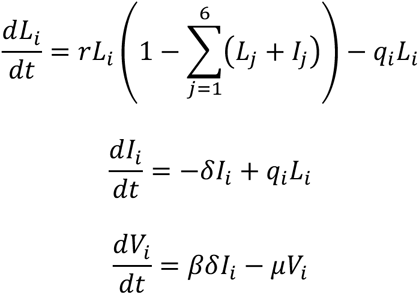

We have partially non-dimensionalized the model so that 1 time unit in the simulations corresponds to about 50 minutes. The lysogeny growth rate is denoted by *r*, the spontaneous induction rates by *q_i_*, *i* = 1, …, 6, the rate of phage production by *δ*, the burst size is by *β*, and the rate of viral degradation by *μ*. The first two equations are decoupled from the last one describing the phage (since we do not consider superinfections in this model). Therefore, the dynamics of the bacterial compartments resemble those of generalized Lotka-Volterra competition.

### Statistical measures

Data visualization and statistical analyses were performed in R version 4.3.1 (R Core Team, 2023) using the packages tidyverse version 2.0.0 (Wickham et al., 2019), car version 3.1-2 (Fox & Weisberg, 2019), rstatix version 0.7.2, and emmeans version 1.8.6.

## Results

### Evolution of a self-targeting lysogen results in decreased spontaneous induction in exponential and stationary phase

After 12 days and ∼72 generations of exponential growth in rich media, we found that the spontaneous induction of DMS3 lysogens was significantly reduced compared to ancestral isolates in exponential phase (Fig 1; ANOVA, *F*_3,68_ = 16.7, *P* < 1e-8). Isolate induction rates within and between experimental replicates ranged from 0.1% to 0.71% of the culture, while the induction rates of ancestral PA14 lysogen (Lys2) isolates ranged from 0.38% to 1.1% of the culture (Fig 1A). Significant differences were robust to the use of other metrics to estimate spontaneous induction (Zong et al., 2010) (see Fig S3). Spontaneous induction was also significantly decreased in stationary phase for all evolved lysogens (Fig S4; ANOVA, *F*_3,109_ = 86.98, *P* < 2.2e-16), although compared to exponential phase, induction was very reduced in stationary phase overall, indicating that the majority of spontaneous induction takes place in exponential phase in DMS3 lysogens.

**Figure 1.**
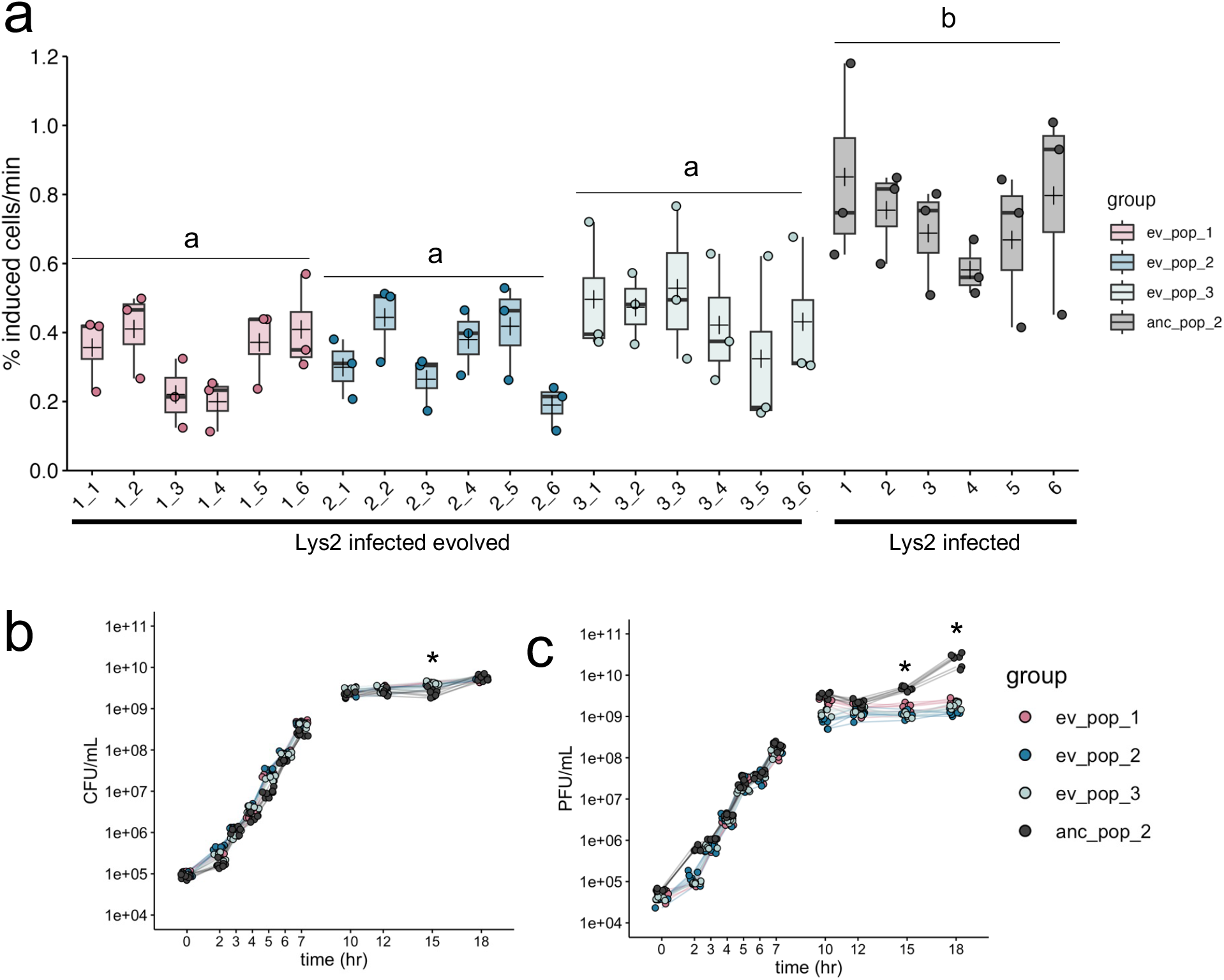
Experimental evolution results in lowered lysogen spontaneous induction. A) Spontaneous induction was measured in exponential phase in six individual isolates from each of three evolved lysogen replicates (ev_pop_1, pink), (ev_pop_2, blue), (ev_pop_3, light blue) and the ancestral strain Lys2 (anc_pop_2, grey). Points are the means of three technical replicates. Bars in the boxplots represent the median; crosses represent the means. Upper and lower bounds of the box are the upper and lower interquartile ranges. Significance was tested with an ANOVA (F_3,68_ = 23.27, p-value = 1.78e-10). Letters indicate significance; groups with different letters have a p-value <0.05; groups with the same or overlapping letters have a p-value of >0.05. B) Bacterial growth curve of all isolates through 18 hours, measured by CFU. C) PFUs sampled through growth curve of all isolates (except the non-lysogenic WT PA14). In B) and C), stationary phase was measured in separate experiments, as indicated by the line breaks between 7 and 10 hours. Points represent the mean of three biological replicates. Asterisks represent a significant difference between the ancestral and evolved populations. Significance was calculated with a Mann-Whitney U test per time point.

### Lysogen populations maintain diversity in CRISPR presence and function

Sequencing of lysogens from each population showed that lysogens evolved the CRISPR locus through a combination of mutation and large deletions and other structural variants (Table 1, Fig 2). These mutations did not overlap with uninfected evolved populations, which exhibited point mutations in flagellar and quorum sensing loci (Fig S1, Table S3), typical of other laboratory evolution experiments (Schick et al., 2022). 39% (7/18) of isolates mutated or deleted the mismatched spacer in the CRISPR2 array, which contains the mismatched spacer which targets the integrated DMS3 (Fig 2B, Fig S5). Two mutations occurred in parallel between two different replicate populations in the evolved lysogen treatment: an A to G point mutation in the seed region of the self-targeting spacer, and an exact deletion of the self-targeting spacer and its upstream repeat (Fig 2B, Fig S6). 22% (4/18) of isolates had disruptions (three independent frameshift mutations and a small deletion) within the *cas* genes *cas7* and *cas8* which form part of the complex that mediates interference in *P. aeruginosa* (Chowdhury et al., 2017). Of these three frameshift mutations, one led to the predicted loss of the *cas7* RNA-binding domain, and two are predicted to interfere with the cas3*-*recruitment or interaction domain of cas8 (Fig 2B, Table 1). One strain was recovered with a 3 kb deletion in the *cas* gene region that spans *cas5* (also part of the interference complex) and *cas7*. While each strain with mutations and indels in the CRISPR array likely no longer target the DMS3 prophage, they maintain CRISPR function. We confirmed that the evolved lysogens did not acquire new spacers (Fig S6).

**Figure 2.**
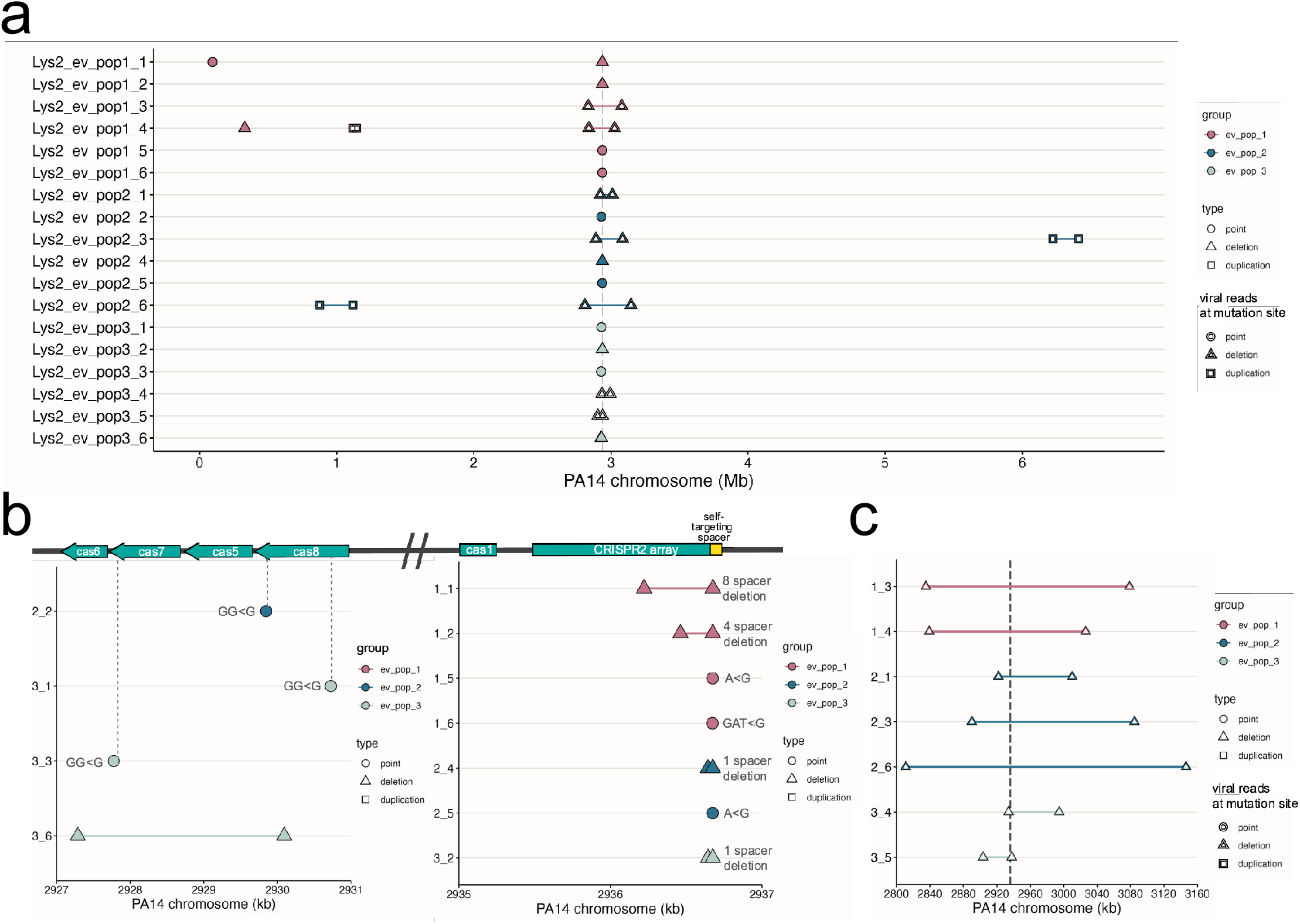
Mutations in infected evolved strains are distinct from uninfected evolved strains. A) Graph of mutations found in all infected evolved samples. B) Close-up of point mutations and small deletions in infected evolved strains. The majority are found in the self-targeting spacer 1 in the 2^nd^ CRISPR array. Gray dashed lines indicate location in the genes. C) Close-up of large deletions in infected evolved strains. Gray dashed double lines indicate the boundaries of the CRISPR-Cas region. In A-C), the y-axis indicates sample ID; x-axis indicates position on the PA14 reference chromosome. Points represent point mutations; triangles spanned by a segment represent deletions of the spanned region; squares spanned by a segment represent duplications of the spanned region. White fill indicates a mutation that was caused by a virus.

**Table 1.** Description of mutations recovered in evolved lysogens.

| Group | Sample ID | Mutated region | Description | Due to phage | Location on PA14-REF chromosome |
| --- | --- | --- | --- | --- | --- |
| ev_pop1 | ev_pop1_1 | CRISPR2 sp1-8 | Deletion | No | 2936222-2936676 |
| ev_pop1 | ev_pop1_1 | *tssK* | Type 6 secretion system baseplate gene; Nonsynonymous | No | 94370 |
| ev_pop1 | ev_pop1_2 | CRISPR2 sp1-4 | Deletion | No | 2936462-2936675 |
| ev_pop1 | ev_pop1_3 | Δ 243,737 bp | Includes CRISPR region | Yes | 2835111-3078848 |
| ev_pop1 | ev_pop1_4 | Δ 186,866 bp | Includes CRISPR region | Yes | 2839227-3026093 |
| ev_pop1 | ev_pop1_4 | 27,769 bp | Duplicated region | Yes | 1120309-1148078 |
| ev_pop1 | ev_pop1_4 | *cysT* | Sulfate transport protein; (CAG)<sub>4→3</sub> | No | 329528 |
| ev_pop1 | ev_pop1_5 | CRISPR2 sp1 | A<G mutation in the 2<sup>nd</sup> nt | No | 2936674 |
| ev_pop1 | ev_pop1_6 | CRISPR2 sp1 | GAT<G deletion of 1<sup>st</sup> and 2<sup>nd</sup> nt | No | 2936673 |
| ev_pop2 | ev_pop2_1 | Δ 88,123 bp | Includes CRISPR region | Yes | 2921860 |
| ev_pop2 | ev_pop2_2 | *cas8* | Frameshift (predicted loss of C-term helical bundle region) | No | 2929846 |
| ev_pop2 | ev_pop2_3 | Δ 194,512 bp | Includes CRISPR region | Yes | 2890191-3084703 |
| ev_pop2 | ev_pop2_3 | 187,901 bp | Duplicated region | Yes | 6222744-6410645 |
| ev_pop2 | ev_pop2_4 | CRISPR2 sp1 | Deletion | No | 2936643-2936675 |
| ev_pop2 | ev_pop2_5 | CRISPR2 sp1 | A<G mutation in the 2<sup>nd</sup> nt | No | 2936674 |
| ev_pop2 | ev_pop2_6 | Δ 335,331 bp | Includes CRISPR region | Yes | 2811042-3146373 |
| ev_pop2 | ev_pop2_6 | 244,019 bp | Duplicated region | Yes | 876284-1120303 |
| ev_pop2 | ev_pop2_6 | Intergenic region | (GCCAAC)<sub>11→8</sub> | No | 3792568 |
| ev_pop3 | ev_pop3_1 | *cas8* | Frameshift (retention of first 57/435 aa) | No | 2930725 |
| ev_pop3 | ev_pop3_2 | CRISPR2 sp1 | Deletion | No | 2936644-2936675 |
| ev_pop3 | ev_pop3_3 | *cas7* | Frameshift (predicted loss of RNA-binding domain) | No | 2927776 |
| ev_pop3 | ev_pop3_4 | Δ 60,538 bp | Begins in *cas3* gene and includes CRISPR2 array | Yes | 2934007-2994541 |
| ev_pop3 | ev_pop3_5 | Δ 34,249 bp | Includes CRISPR region | Yes | 2903549-2937794 |
| ev_pop3 | ev_pop3_6 | Cas gene deletion | Deletion of *cas7* and *cas5*, partial deletion of *cas6* and *cas8* | No | 2927288-2930093 |

We observed that 44% (8/18) of evolved lysogen isolates carried deletions of varying sizes; the smallest being the 3 kb deletion in the *cas* genes, and the largest deletion, 335,331 bp, comprising about 5% of the PA14 chromosome. Of these eight deletions, seven were centered on the CRISPR2 array (Fig 2C). Therefore, in the evolved lysogen populations, we observed extensive coexistence between combinations of CRISPR spacer mutations and large entire deletions of CRISPR, demonstrating the importance of phage infection in determining distinct evolutionary trajectories in isolates in the same environment.

### Evolved lysogens with large deletions are polylysogens

While confirming that the evolved lysogens had retained the phage at its original integration site (Methods), we found that the boundaries of the deletions of the CRISPR regions were composed of reads that mapped to both the PA14 and DMS3 chromosomes (hereafter referred to as “split” reads), indicating that the large deletions in these seven isolates resulted from a DMS3 transposition event which occurred from within the chromosome (Fig 2C). Therefore, we consider these deletions to be phage-mediated. These phage-mediated deletions ranged from approximately 34-335 kb (mean = 163.3 kb ± 100.3 kb). Deletions occurred in each of the replicates independently, with no shared deletion boundaries even within the same culture (Fig 2C). The regions of the phage chromosome to which the split reads mapped and the orientation of phage reads at the boundaries of the deletions in two samples (1_4 and 2_3) suggests that more than one phage genome may be inserted in the gap (Fig 3D-E, Table 2). Notably, we did not recover any mutations in the phage chromosome.

**Figure 3.**
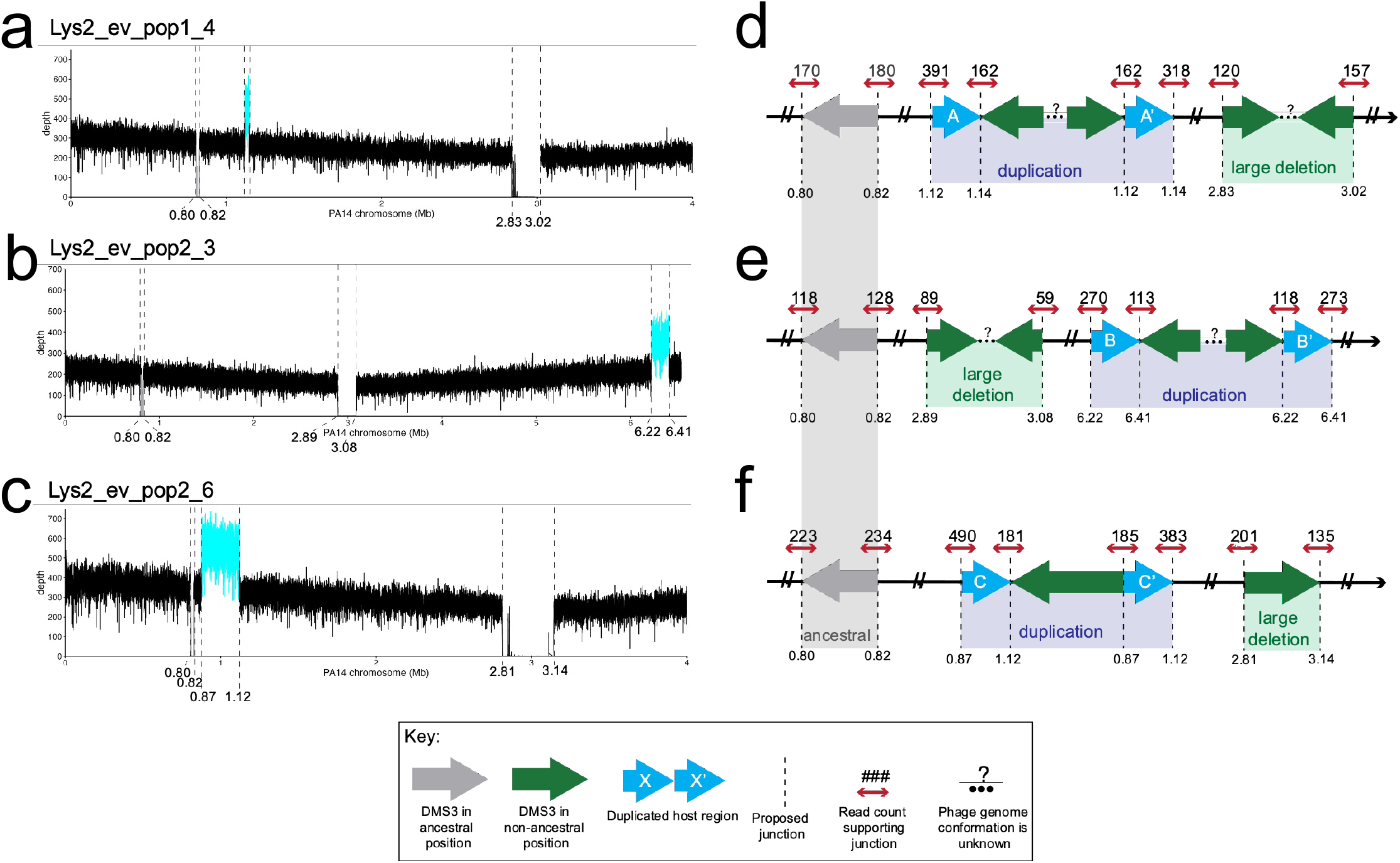
Location and characterization of large duplicated regions in three evolved lysogen isolates. A-C) Coverage plots of the genome. Dashed lines indicate evidence of host-phage boundary. X-axis is position on the PA14 chromosome. Where possible, the x-axis has been truncated to zoom on all possible insertion sites. Y-axis is depth of coverage at that nucleotide. Cyan line represents the putative duplicated region in between two viral insertion sites. Gray line is the region between the ancestral insertion sites. In D-F) Cartoon of resulting genome architecture of the evolved lysogens. Dashed lines indicate a putative host-phage boundary. Gray arrows represent the phage at its ancestral insertion site; green arrows represent phage at new insertion sites. Blue arrows represent duplicate host sequence. The direction of the arrow indicates 5’ to 3’, with 3’ ending at the tip of the arrowhead. Dashed lines represent the junctions between phage and host. Small red arrows indicate host reads; numbers above the arrow indicate read count at that site that supports that junction. Ellipses and question marks between phage genomes represent uncertainty. Read files for each sample and at each location can be found in Supplementary Data. In A and C, regions of coverage in the deletion region in Lys2_ev_pop1_4 and Lys2_ev_pop2_6 are from domains in a TpsA1 and TpsB1 protein (annotated as a filamentous hemagglutinin protein), and in the 3’ end of the Lys2_ev_pop2_6 deletion, a domain in OprB (annotated as a porin). Both were identified via BLASTX.

**Table 2.** Origin of phage reads at the deletion boundaries. Asterisks denote samples where reads from only one end of the phage genome were recovered.

| Sample ID | Group | <u>Upstream deletion boundary</u><br>Read mate maps to: | <u>Downstream deletion boundary</u><br>Read mate maps to: |
| --- | --- | --- | --- |
| Lys2_ev_pop1_3 | ev_pop_1 | end of DMS3 genome | start of DMS3 genome |
| *Lys2_ev_pop1_4 | ev_pop_1 | start of DMS3 genome | start of DMS3 genome |
| Lys2_ev_pop2_1 | ev_pop_2 | end of DMS3 genome | start of DMS3 genome |
| *Lys2_ev_pop2_3 | ev_pop_2 | start of DMS3 genome | start of DMS3 genome |
| Lys2_ev_pop2_6 | ev_pop_2 | end of DMS3 genome | start of DMS3 genome |
| Lys2_ev_pop3_4 | ev_pop_3 | end of DMS3 genome | start of DMS3 genome |
| Lys2_ev_pop3_5 | ev_pop_3 | end of DMS3 genome | start of DMS3 genome |

The large deletions, though centered around the CRISPR locus, had different deletion boundaries. To assess the shared gene content in these regions, we used eggNOG-mapper to query the functional protein content that was lost (Methods). In addition to CRISPR, the deleted regions were enriched with genes from COG category S (genes of unknown function), which included Type 6 secretion system-related genes *tssF* and *tssG*, and multiple pyoverdine system genes which are often lost during lung colonization (Nguyen et al., 2014; O’Brien et al., 2017; Schick & Kassen, 2018). Several pyoverdine biosynthesis and transport genes (*fpvA, pvdE, pvdH, pvdI, pvdM, pvdM, pvdO*) were deleted in 6 of the 7 isolates with large deletions. *PvdS,* a regulator of pyoverdine biosynthesis genes (Leoni et al., 2000), and *pvdR,* a component of the pyoverdine efflux transporter (Imperi et al., 2009), were also deleted in 5 of the 7 isolates with large deletions. The co-localization of these virulence and defense gene cassettes with CRISPR may contribute to variation in these regions (Makarova et al., 2011; L. A. Robinson et al., 2023).

In this study, 100% (7/7) of isolates with large deletions were polylysogens; however, polylysogeny did not evolve in cells that resolved self-targeting via CRISPR mutations from SNPs or indels (Fig S7). Similarly, a previous study showed that PA14, when challenged with free DMS3 virions and subjected to a short-term evolution experiment, evolved genome deletions encompassing the CRISPR region when the susceptible host did not have self-targeting (e.g. contained a *cas7* deletion) (see: Extended Data Table 1 in Rollie *et al*, 2020). In combination with our data, we infer that self-targeting increases the rate of DMS3 transposition, resulting in polylysogens, which then are maintained in the population only when they are associated with adaptive mutations, such as those which decrease spontaneous induction by resolving genetic conflict (Fig S7; for model, see Fig 6).

### Evolved polylysogens contain large duplications

Three isolates containing phage-mediated large deletions in CRISPR also had large duplications elsewhere in the genome (27, 188, and 244 kb; mean = 153 ± 113 kb). In these cases, these regions were not deleted but were doubled in coverage (Fig 3A-C). The location of these large duplications exactly corresponded to additional insertion sites which were recovered by our pipeline (Methods). Due to the orientation of the PA14/DMS3 split reads at these insertion sites, which faced away from each other rather than toward each other, and due to the fact that the split reads at the boundaries of the duplicated regions only represented about 50% of the total coverage, we interpret these regions to be large duplications with a phage genome in the middle, as opposed to two independent viral insertions (Fig 3D-F). As the duplicated regions were not centered around a shared core, they were almost completely non-overlapping in their gene content. Only one known gene was duplicated in two of the three isolates (Lys2_ev_pop2_3 and Lys2_ev_pop2_6), which was *betT,* a choline transporter known to accumulate mutations in clinical isolates from CF patients (Malek et al., 2011; Marvig et al., 2015; Stribling et al., 2023). Broadly, genes from category H (coenzyme metabolism) were represented in all three isolates, as well as from P (inorganic ion metabolism) and S (genes of unknown function), as in the large deletions. Several genes annotated as part of the major facilitator superfamily, a class of membrane-associated transporter proteins, were also duplicated in two of the three isolates, as well as genes from the *moa* family, which have recently been implicated in biofilm formation (Kaleta & Sauer, 2023).

Two of these duplications (in 1_4 and 2_6) independently evolved a shared boundary six nucleotides apart (at positions 1120309 and 1120303, respectively), in an intergenic region between the 3’ end of a hypothetical protein and the 3’ end of an AraC transcriptional regulator. An analysis of all new lysogen insertion sites (including the deletion and duplication boundaries) using the motif-finding software MEME Suite did not return any motifs, either using MEME (searching for a motif in a 15 bp region centered on the insertion site), or MEME-ChIP (searching for centrally enriched motifs 250 bp around the insertion site) (Bailey & Elkan, 1994; Machanick & Bailey, 2011); this suggests parallel duplications may be advantageous in this environment.

Neither the isolates containing large deletions nor the ones containing large duplications differed in their growth from other evolved strains that did not have large structural variation (Fig S8A). This suggests that, under these conditions, the fitness costs to deletions, duplications or carrying additional copies of the phage in the chromosome are smaller than the fitness gains by removing self-targeting. Although these phage-mediated deletions of CRISPR represent the addition of one to two phage genomes to the lysogen chromosome, the spontaneous induction rate of these isolates remains reduced relative to the ancestral PA14 lysogen strain (Lys2) (Fig 1, Fig S8B). Additionally, although viral output does not change with phage genome copy number after challenge with mitomycin C, cell survival is significantly increased with increased phage genomes (Fig S8C), suggesting a possible mechanism of viral interference leading to cell survival which may also contribute to a decreased spontaneous induction measurement.

### Spontaneous induction correlates with mutation

We observed that spontaneous induction was variable within replicate populations (Fig 1). We asked whether this variation might correlate with differences in the type of CRISPR mutation (SNP, deletions, viral transpositions). To address this, we grouped lysogens into 1 of 5 categories based on the type of mutation that occurred in the genome (“cas deletion”; “cas mutation”; “spacer deletion”; “spacer mutation”; and “large deletion polylysogen”) and asked whether including mutation type reduced the variation within these groups. We observed that all groups (with the exception of the cas deletion group, which had only one isolate in its group) had significantly lower spontaneous induction than the ancestral strains (Fig 4); and the large deletion polylysogen group was significantly lower than lysogens that had lowered spontaneous induction via SNPs or indels. Another set of experiments which included a lysogenized ΔCRISPR strain showed that evolved lysogens which resolved genetic conflict via genome rearrangements, SNPs, and indels, reduced CRISPR function to the level of a ΔCRISPR mutant (Fig S9). A model incorporating mutation type was a significantly better fit than the model by experimental replicate (Fig 4, ANOVA, *F*_2,66_ = 13.777, *P* < 1e-6). In view of these results, we find that heterogeneity in genotype correlates to the heterogeneity in phenotype of spontaneous induction in our evolved lysogens.

**Figure 4.**
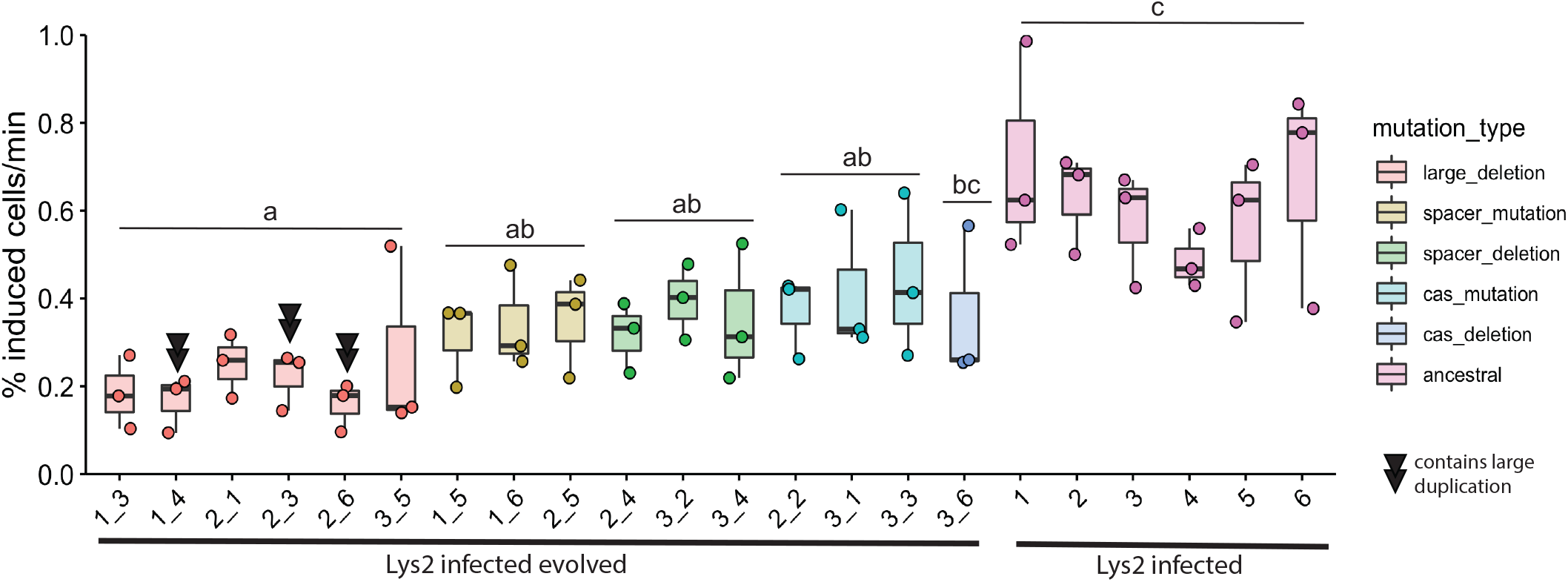
Genomic variation may explain phenotypic variation. Evolved isolates were grouped by type of mutation. Colors of points and boxes represent groups. Points are means of experimental replicates. Box plots are as in Figure 1. Isolates with double black triangles have large duplications which are due to another viral insertion. Significance was determined with an ANOVA with Tukey adjustment using the log-transformed values (F_5,66_ = 19.3, p-value = 8.725e-12). Letters indicate significance; groups with different letters have a p-value <0.05 between them; groups with the same or overlapping letters have a p-value of >0.05 between them.

Given these small but significant variations in spontaneous induction that are maintained within groups yet replaced the ancestral lysogen genotype, we wanted to understand how long the diversity we observed within our experimental replicates could persist in exponentially growing cultures. To do this, we developed a mathematical model to compare six lysogens with six different rates of spontaneous induction (two values from the ancestral group, two values from the host-mutation group, two values from the large deletion polylysogen group). In a deterministic model of lysogen growth in exponential phase with varied spontaneous induction rates, we founded populations with low densities of high inducers (representing the ancestor strain) and allowed them to grow for 10 hours. At that time, we introduced either strains with low rates of spontaneous induction (representing the large deletions) or medium rates of spontaneous induction (representing host mutations), one at 10 hours and the other at 24 hours, and tested whether they could invade. We found that when low inducers are introduced to a high-inducing population, medium inducers cannot subsequently invade (Fig 5A). When we introduced medium inducers to a high-inducer population after 10 hours of growth, and then low inducers after 24 hours, medium and low inducers outcompeted the high inducers and then coexisted in the absence of the high inducers for about two days (Fig 5B). This coexistence recapitulates the observed recovery of low and medium inducers, but not high inducers, in our experimental data (Fig 4). From these observations we find that the weak selection imposed by these small differences in spontaneous induction, which are caused by different mutational mechanisms, combined with the order in which they were introduced, may allow evolution of diversity in CRISPR self-targeting resolution in *P. aeruginosa* lysogens and preserve coexistence of CRISPR+ and CRISPR-strains in the absence of additional selective variables or environmental change.

**Figure 5.**
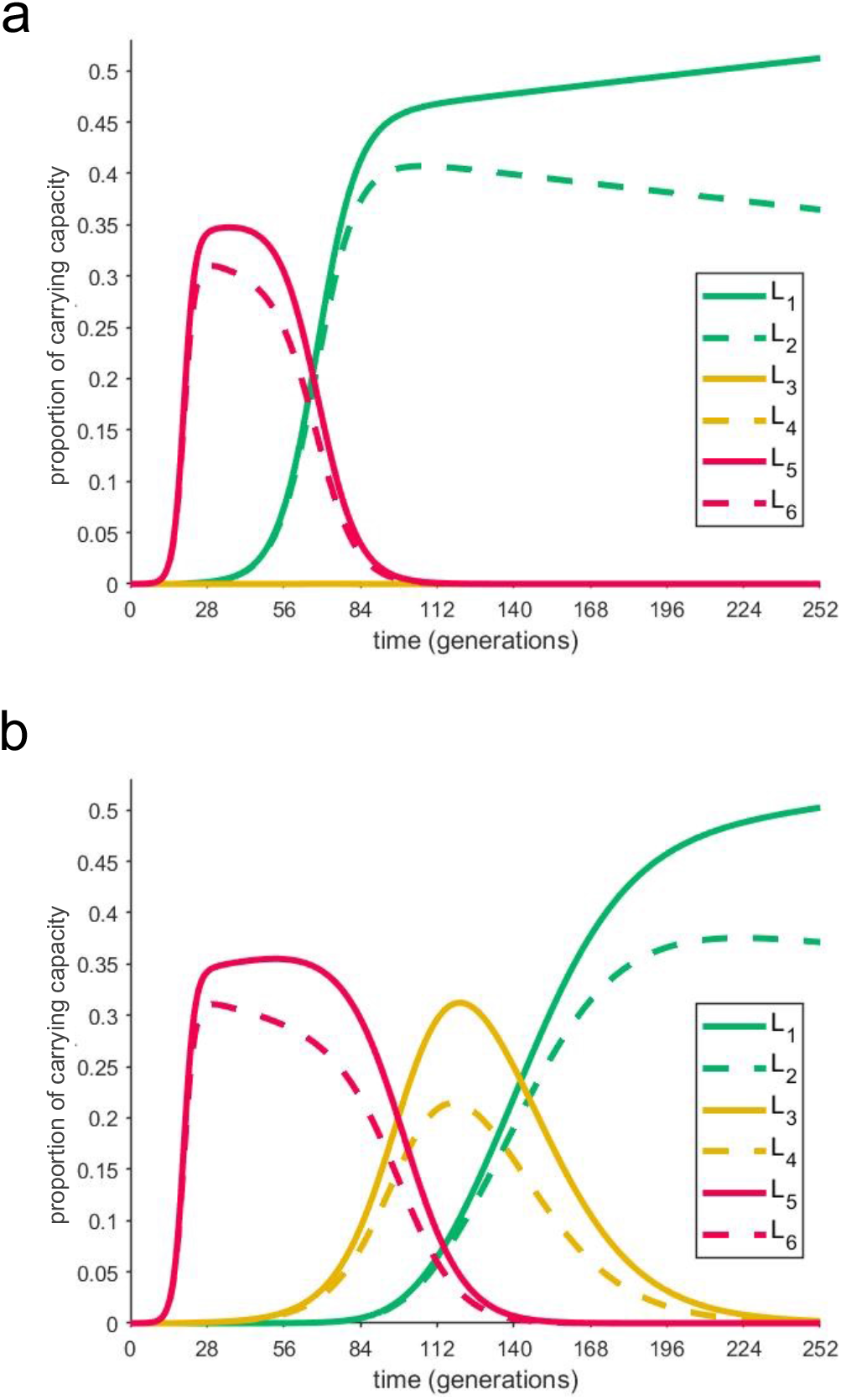
Coexistence of different rates of spontaneous induction depends on the order of their introduction. A mathematical model describes the behavior of several lysogens with different spontaneous induction values. These graphs describe the growth of six lysogens over time in one continuously growing culture. The y-axis represents the proportion of carrying capacity of the medium; the x-axis is time in PA14 generations. Each lysogen was assigned a spontaneous induction value from the experimentally determined range. Each pair (L1/L2; L3/L4; L5/L6) in a certain range are taken from values in the same group, from lowest to highest spontaneous induction values (L1 is the lowest; L6 is the highest). In A) the highest spontaneous inducers are introduced first. At approximately 9 hours, low inducers are introduced; after about 1 day, medium inducers are introduced and do not establish. In B) the highest spontaneous inducers are introduced first. At approximately 9 hours, medium inducers are introduced; after about 1 day, low inducers are introduced and establish, resulting in a period of coexistence between medium and low inducers that is observed experimentally.

**Figure 6.**
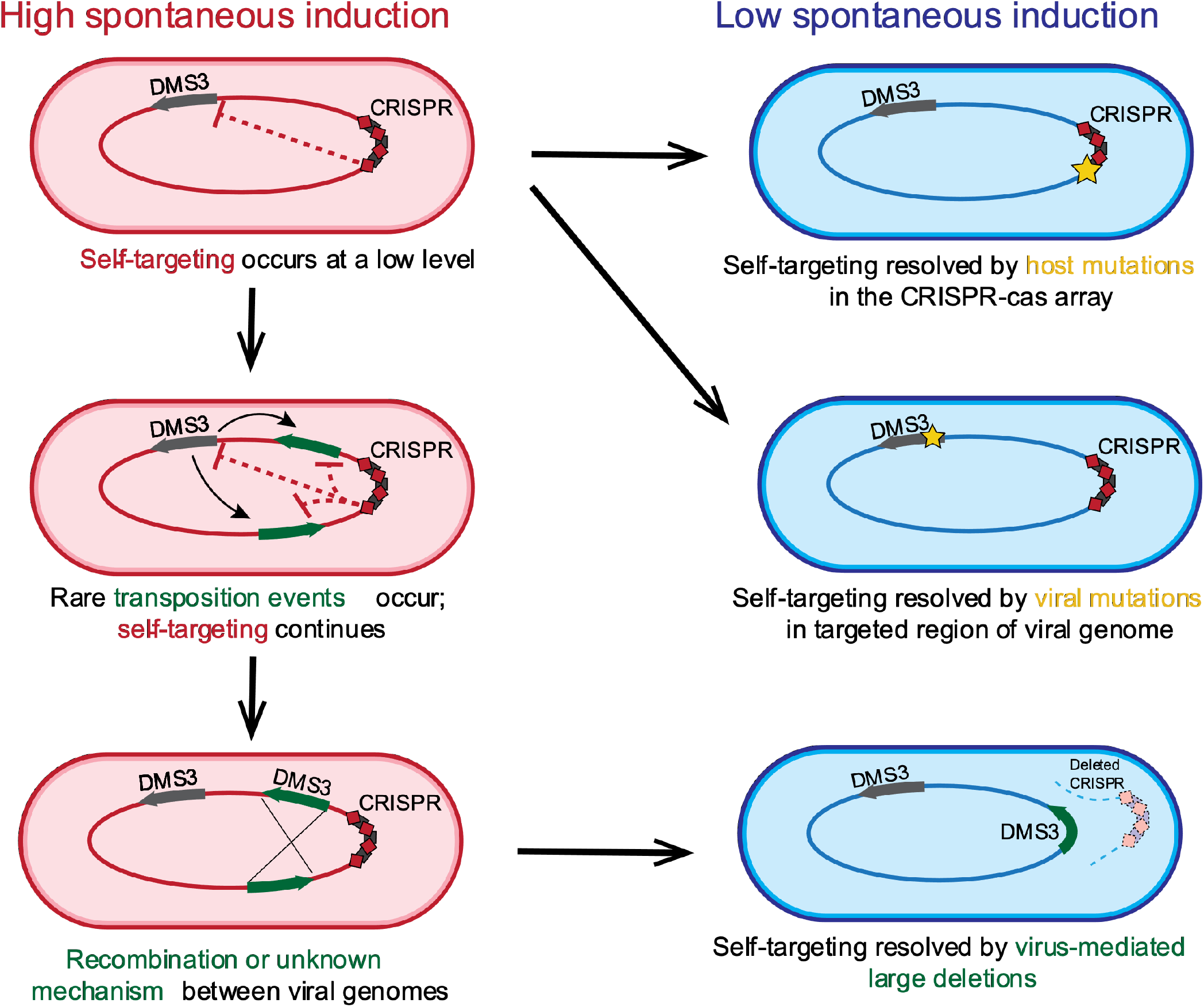
Model of self-targeting leading to DMS3 transposition around the genome. Spontaneous induction caused by CRISPR self-targeting is resolved in two modes. Cells with high spontaneous induction are indicated in the first column, in red. One mode, in the first column, relies on low levels of DMS3 escape from lysogenic repression which result in rare transposition events around the genome. Recombination between multiple DMS3 chromosomes leads to large deletions, which may include the CRISPR area and result in cells with lower spontaneous induction (blue). In the second mode, self-targeting may be resolved by host mutations in the CRISPR-Cas region, or by viral mutations in the targeted region (not recovered in this study).

## Discussion

Although temperate phages are frequently recovered from long-term, evolving *Pseudomonas* infections of the lungs of CF patients, phages are often considered at single time points or outside of relationship to their bacterial hosts (Ambroa et al., 2020; Gabrielaite et al., 2020; Tariq et al., 2015, 2019). Evolution experiments involving transposable phages often include susceptible cells, or are begun with free phages instead of established lysogens (Davies et al., 2016; Marshall et al., 2021; O’Brien et al., 2019; Rollie et al., 2020). Because these transposable phages may insert into the bacterial chromosome in many different locations, this approach can unite selection on the phage with the production of beneficial bacterial mutations that are generated upon infection. Here, we bypass this issue to query the impact of a phage on the evolution of its host genome by studying the context of an established lysogen (vertical transmission), and find that the presence of temperate phage alone profoundly changed the genome of the host through genomic rearrangements mediated by polylysogeny from within. This is especially relevant to understanding *P. aeruginosa* infections which may persist for years in the absence of phage lysis and horizontal transmission.

Our work supports previous studies showing that DMS3 lysogens evolve decreased rates of spontaneous induction over time (Rollie et al., 2020; Zegans et al., 2009); here, we distinguish between exponential and stationary phase, and find that exponential phase induction accounts for the majority of free phages in the medium. Our spontaneous induction estimates in exponential phase are approximately 0.37% and 0.72% of the evolved and ancestral populations, respectively, which are both significantly higher than lambda-like phages (Imamovic et al., 2016; Little et al., 1999; Zong et al., 2010), but are in line with other studies that report high spontaneous induction (Owen et al., 2017). Mu-like phages have been reported previously to naturally produce high titers during lysogen growth (Bondy-Denomy et al., 2016; James et al., 2012; Zegans et al., 2009), suggesting that this evolutionary pressure is not restricted to DMS3 and Mu-like phages as a family have high spontaneous induction rates. Because Mu-like phages are both highly prevalent and highly targeted in *P. aeruginosa* (England et al., 2018), it is likely that CRISPR self-targeting will inform evolutionary outcomes in these related phages. Here, the resolution of that self-targeting resulted in coexisting variation in lysogen spontaneous induction rates in exponential growth, with polylysogens having the lowest rates perhaps due to viral interference preventing cell death, and lowering virion production (Refardt, 2011). These lysogens could also have lower rates not because of polylysogeny, but because of the large deletion itself.

We find that the genetic conflict between Mu-like phage and host results in a tradeoff between CRISPR immunity and spontaneous induction, which could help explain the maintenance of CRISPR systems in *P. aeruginosa* (Cady et al., 2011; Soliman et al., 2022; Van Belkum et al., 2015). Previously, mutations in *cas7* (Zegans et al., 2009), and deletions of the entire CRISPR region (Rollie et al., 2020) have been found to reduce phage induction in late log phase. We find that these differences in self-targeting resolution reduce phage induction to different degrees and have profoundly different effects on the host genome. Half of the evolved lysogen isolates decreased their spontaneous induction while maintaining either CRISPR function (in mutations or deletions of the self-targeting spacer) or the potential of CRISPR function (frameshift mutations in *cas* genes), while the rest deleted CRISPR and lost any potential immunity, but more substantially decreased their induction. One explanation for the maintenance of these two genotypes is the order in which these mutations were introduced. Modeling simulations show that spontaneous induction rates from strains which resolved self-targeting via SNPs and indels (“medium” inducers) cannot invade established populations of strains with spontaneous induction rates from polylysogens. This indicates that host-mediated SNPs and indels likely arose before polylysogeny and large deletions, as both are maintained together despite spontaneous induction differences. This suggestion of an order (host-mediated before phage-mediated) then suggests that the basal rate of phage transposition is lower than host mutation. A lower rate of formation of higher-fitness polylysogen “low” inducers, which compete with lower-fitness host-mutation “medium” inducers with a faster rate of formation, may work to maintain a pool of diversity that selection may subsequently act on in different ways, given the presence of other phages or other ecological factors (Watson et al., 2023).

In an isolated environment, our model predicts that diversity would likely be resolved by the eventual replacement of CRISPR-immune cells with non-immune polylysogens, thereby increasing the phage copy number in an environment that does not have new susceptible hosts for horizontal transmission (Weitz et al., 2019). However, complex communities with additional phage and bacteria may then select for cells which maintain CRISPR function but which have higher spontaneous induction (Alseth et al., 2019; Gloag et al., 2019). A limitation of this study is that it does not take into account the highly polymicrobial nature of the cystic fibrosis lung and uses only one phage-host pair to explore the evolutionary outcomes of lysogeny. These outcomes may be specific and vary with the evolutionary history of each phage-host pair. How phages evolve increased fitness while maintaining CRISPR immunity and defense against other phages and bacterial competitors in complex environments remains an open question.

The results of genetic conflict in evolved lysogens are not limited to CRISPR deletions and may impact the rate of evolution of bacteria with latent infections. Gene loss and genome reduction have also been shown to occur in *P. aeruginosa* lineages during adaptation to the human lung, although the contribution of phages to this loss is unclear (Gabrielaite et al., 2020; Rau et al., 2012). In this study, pyoverdine and Type 6 secretion system genes were lost in the majority of the polylysogens, and one evolved lysogen isolate (1_1) had a nonsynonymous mutation in the Type 6 secretion system baseplate gene *tssK* (Table 1). These genes, lost under laboratory conditions, are also often lost in chronic CF isolates (Marvig et al., 2015; O’Brien et al., 2017; Perault et al., 2020).

The mechanism of this gene loss is unclear in two isolates. In many evolved polylysogens, a phage genome simply replaced the deleted sequence as in (Rollie et al., 2020). In these isolates, the phage genome appeared in an head-tail or tail-head configuration, which could occur as a result of the replicative transposition reaction itself, or as a result of recombination between two preexisting phages. Recombination between a duplicated sequence is an attractive hypothesis to explain the deletions because it may lead to deletion or duplication of the intervening sequence (Reams & Roth, 2015), and because Cas3 targeting of the phage chromosome may stimulate this recombination. However, two of three isolates with a duplication (1_4 and 2_3) exhibit both non-canonical deletion and duplication structures, where we recover host-phage junctions which suggest two phage genomes facing either head-head (reads recovered at both junctions which map to the 3’ end of the genome) or tail-tail (reads recovered at both junctions which map to the 5’ end of the phage genome). Additionally, recombination may result in two phage-host junctions on the 3’ end of the duplication which lead into different ends of the phage chromosome, whereas we only recover reads which lead into one end of the phage chromosome (Reams et al., 2012). Whether and how phage presence continues to alter the evolution of its host from the uninfected state, and how phage infection influences the rate and mechanism of this evolution, are questions which require future study to explain the ubiquity of phage infection in many clinical environments (Burgener et al., 2019; Holloway et al., 1960; Vaca-Pacheco et al., 1999).

*P. aeruginosa* evolution in the context of the CF lung can occur via a slow accumulation of SNPs and indels (Marvig et al., 2015) or a more rapid accumulation of SNPs due to the evolution of hypermutator genotypes (Marvig et al., 2015; Oliver et al., 2000). Lysogenization by transposable phages may offer a different mechanism of within-lung diversification which operates in addition to the baseline mutation rate. Due to the nature of short-read sequencing, it is likely that polylysogeny of the same virus, and resulting genome rearrangements, have been under-represented in current datasets of *P. aeruginosa* clinical isolates. Future studies should continue to identify signatures of multiple phage infection in clinical isolates, and look for deletions and duplications that may be associated with phages. The inferred dependence of transposition on the presence of self-targeting, and by extension on low levels of DNA damage, also opens the possibility of other causes of low levels of DNA damage – for example, subinhibitory concentrations of DNA-damaging antibiotics – to be driving evolutionary adaptation and diversity in bacteria lysogenized by transposable phages. The extent to which phage infection informs differences in evolutionary and functional outcomes in a clinical context is an important subject for future work.

## Acknowledgements

We thank Santiago Elena at I2SysBio and the University of Valencia for help conceptualizing and establishing initial experimental evolution studies on lysogen fitness. We are grateful to George O’Toole for discussions as well as strains of bacteria and phages described herein including Lys2 and DMS3. We gratefully acknowledge Dr. Alan Collins for helpful discussions during early stages of this project. We thank Whitaker lab members Jiayue Yang for helpful discussions, and Sierra Bedwell and Izzy Lakis for comments on the manuscript. We thank Alvaro Hernandez and Chris Wright of the Roy J. Carver Biotechnology Institute for sequencing expertise, as well as Jeff Haas of the School of Integrative Biology for crucial help with data storage and server access. This work is funded by grants from the Cystic Fibrosis Foundation and the Allen Distinguished Investigator Award from the Paul G. Allen Foundation to R.J.W. and the National Science Foundation grant DMS-1815764 to Z.R. This research is also a contribution of the GEMS Biology Integration Institute, funded by the National Science Foundation DBI Integration Institutes Program, Award #2022049.

## Author Contributions

Conceptualization: L.C.S., R.J.W. Formal analysis: L.C.S. Funding acquisition: L.C.S., R.J.W. Investigation: L.C.S., Z.R., A.C.S. Methodology: L.C.S., Z.R., J.H.C., R.J.W. Resources: J.C.V., R.J.W. Software: J.C.V. Visualization: L.C.S., Z.R. Writing – original draft: L.C.S., R.J.W. Writing – review and editing: L.C.S., R.J.W.

## Supplemental Information

The following parameter values were used in the model equations:

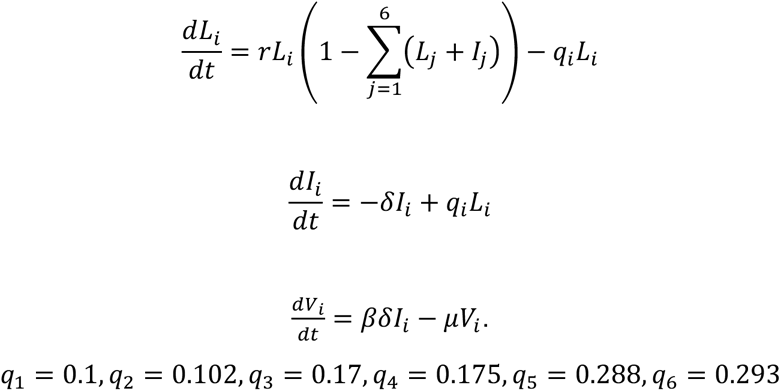

and

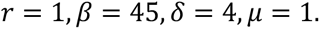

Each generation lasts 38 minutes. The initial conditions used were 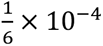 for each *L_i_*, which was added at the prescribed times. *I_i_* and *V_i_* were initially zero for each *i* = 1, …, 6.

**Figure S1.**
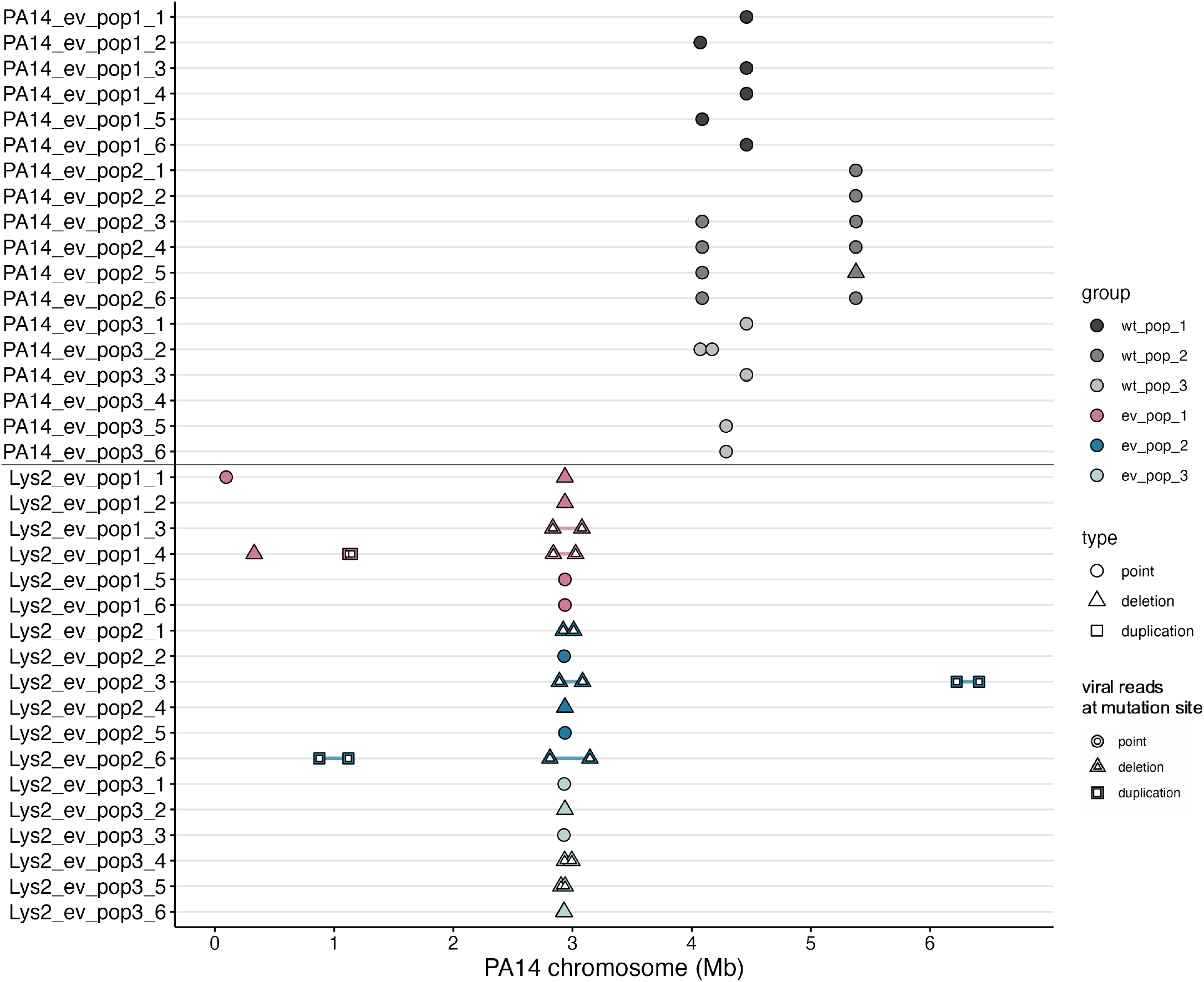
Uninfected populations do not share mutations with lysogen populations. The Y-axis indicates sample ID; X-axis indicates position on the PA14 reference chromosome. Circles represent point mutations; triangles spanned by a segment represent deletions of the spanned region; squares spanned by a segment represent duplications of the spanned region. Inset white shapes indicate a mutation that was caused by a virus. Uninfected population PA14_ev_pop2 contains evidence of phage selection pressure as 100% of isolates have a mutation in the Type 4 pilus. Lysogen data is the same as in Figure 2.

**Figure S2.**
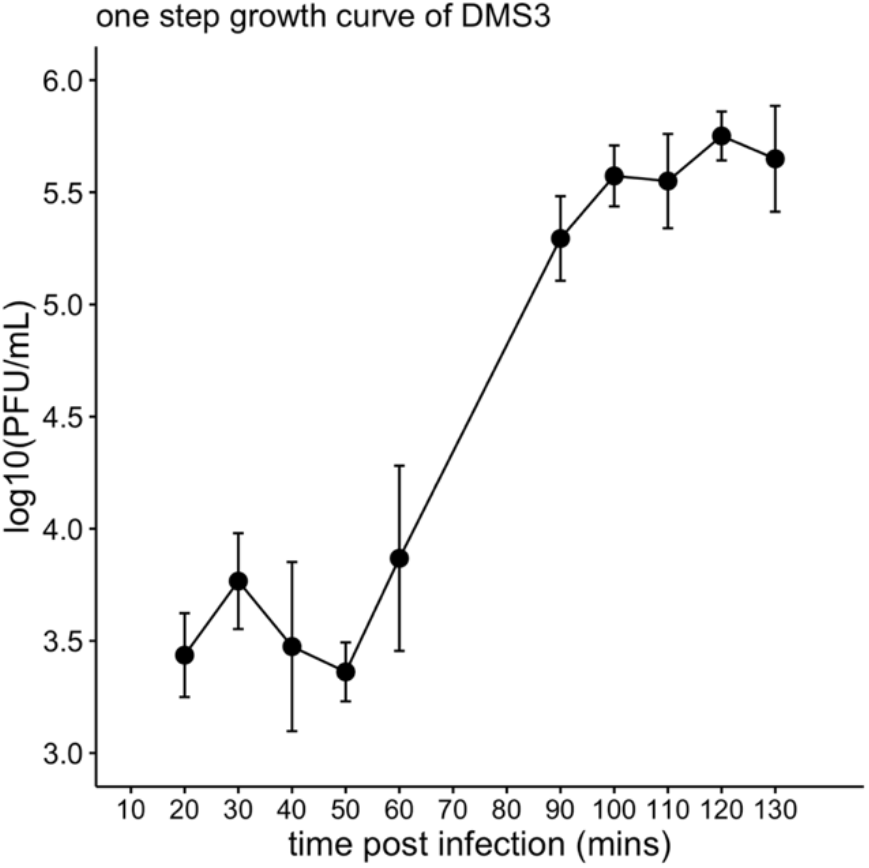
One step growth curve of DMS3. Points represent the mean of experimental replicates (n=6 for all other time points except 20 mins (n=3) and 100 mins (n = 2). Error bars represent standard deviation. The x-axis is time post-infection in minutes; the y-axis is the log_10_ of PFU/mL. Plateau 1 is the average of all values from 20-50 minutes; Plateau 2 is the average of all values from 100-130 minutes.

**Figure S3.**
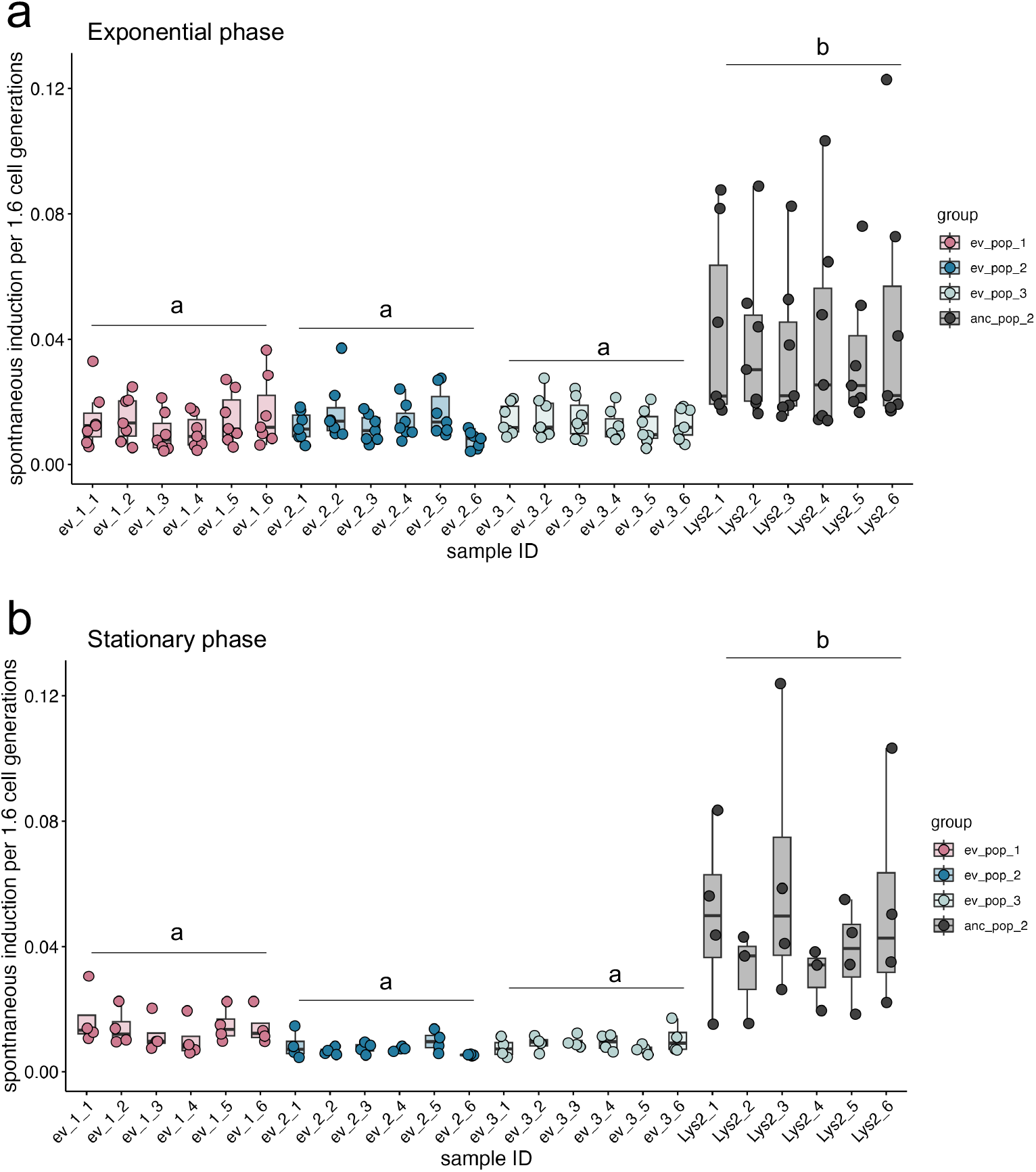
Spontaneous induction calculated with Zong et al formula. A) Spontaneous induction in exponential phase. X-axis is time in hours; y-axis is spontaneous induction per hour, or 1.6 cell generations of PA14. B) Spontaneous induction in stationary phase. X-axis is sample ID; y-axis is spontaneous induction per hour. Significance was calculated as the spontaneous induction value as a function of group with Dunn’s Test of Multiple Comparisons and Bonferroni correction. Letters indicate significance; groups with different letters have a p-value <0.05 between them; groups with the same or overlapping letters have a p-value of >0.05 between them.

**Figure S4.**
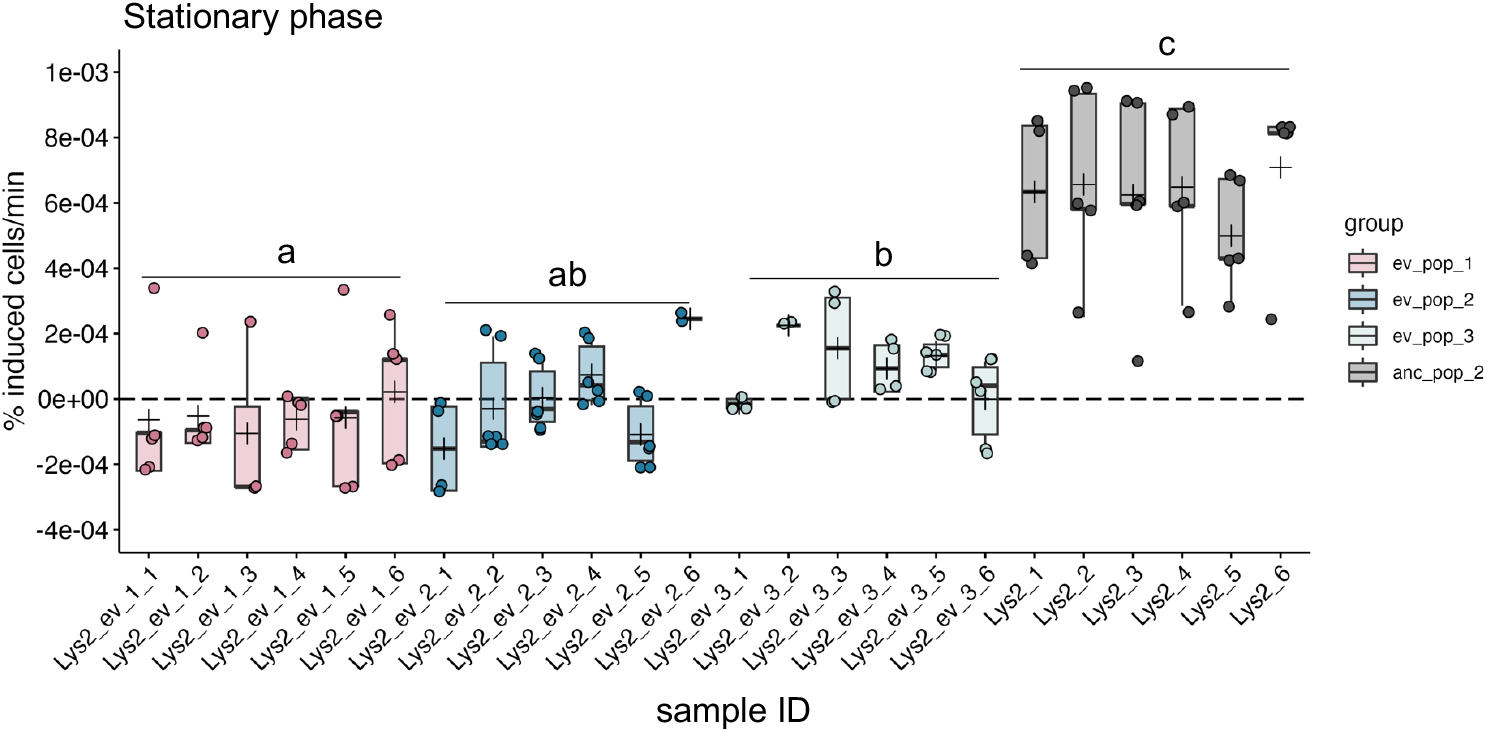
Experimental evolution results in lowered lysogen spontaneous induction in stationary phase. Spontaneous induction was measured in stationary phase. Six individual isolates from each of three lysogen replicates (ev_pop_1, pink), (ev_pop_2, blue), (ev_pop_3, light blue) and the ancestral strain Lys2 (anc_pop_2, grey) were measured. Points are the means of one experimental replicate from three technical replicates. Boxplots are as in Figure 1. Significance was tested with an ANOVA (*F*_3,109_ = 86.98, *P* < 2.2e-16). Letters indicate significance; groups with different letters have a p-value <0.05; groups with the same or overlapping letters have a p-value of > 0.05.

**Figure S5.**
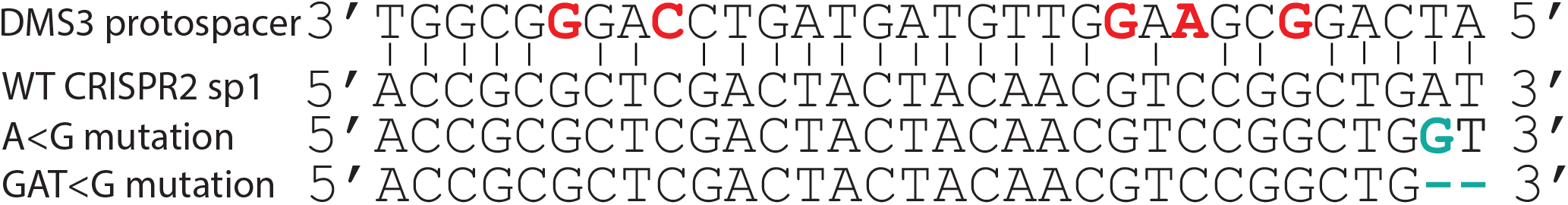
DMS3 is partially matched by CRISPR spacers in PA14 and is resolved by host mutations. Map of spacer-protospacer match compared to escape mutations in the host spacer that evolved in two single isolates in parallel cultures. Five mismatched bases on the virus are highlighted in bold red font and lack lines indicating base pairing; escape mutations are highlighted in blue and lack lines indicating base pairing. Hyphens indicate a deletion of those bases.

**Figure S6.**
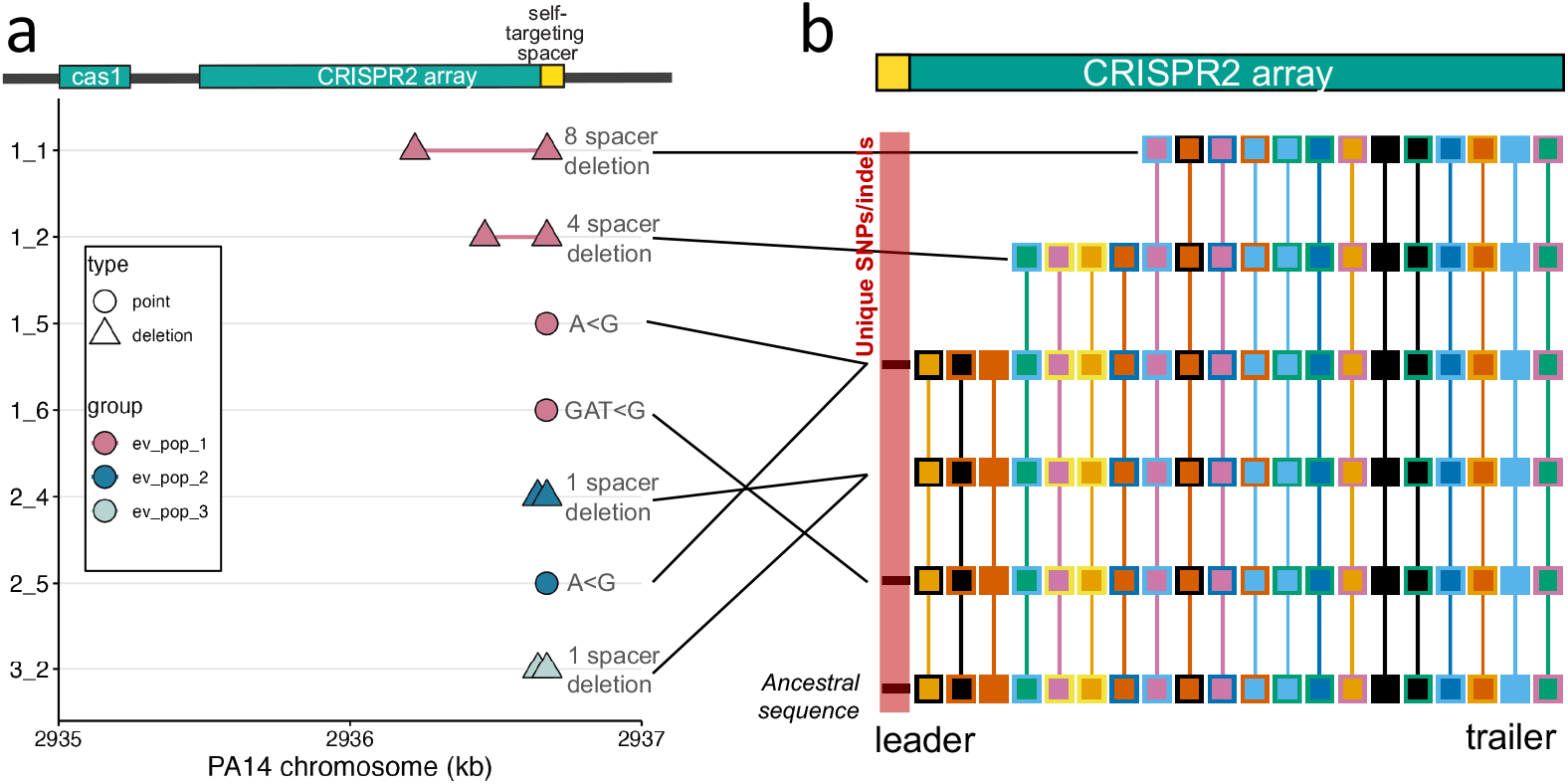
No new spacer acquisition, only spacer evolution in the CRISPR2 array after lysogen evolution. We ran CCTK on the evolved lysogen genomes to look for spacer acquisiton. CCTK recovered all known spacers. A), same figure and data as Figure 2C; B) a CCTK graphical output representing all different arrays from these genomes. Spacers shared between arrays are denoted by boxes with the same colored outline and fill. Spacers that are unique only to that array and which are not shared with any other array are represented as a black line (highlighted with a red box). In our case, these are due not to spacer acquisition but to point mutation which make them non-identical and “unique” with respect to the others. Three cases of unique spacers are due the representation of the ancestral spacer (maintained in all strains which resolve self-targeting via the cas genes) and two unique spacer genotypes.

**Fig S7.**
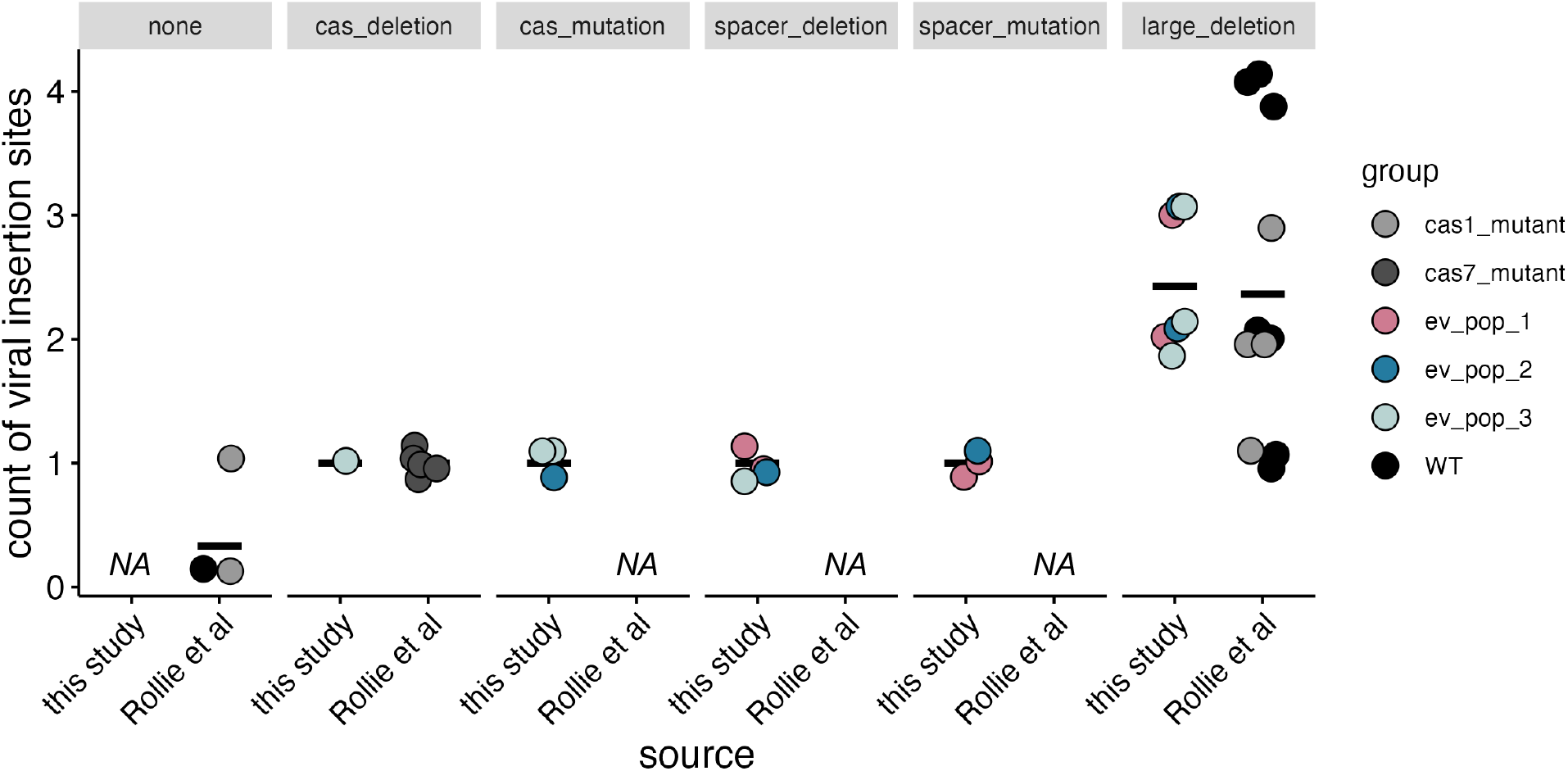
Polylysogeny does not arise in the absence of self-targeting. Y-axis describes the number of viral insertion sites recovered from each isolate; X-axis describes the source of the isolates (either this study or Rollie et al) and is broken up by the type of mutation which the isolate is classified by. Black bars indicate the mean. Points represent one isolate. Point color indicates either what replicate it came from (as in this study) or in what strain background the evolution experiment was started (in Rollie et al). NA: no isolates were recovered in that mutation type category.

**Figure S8.**
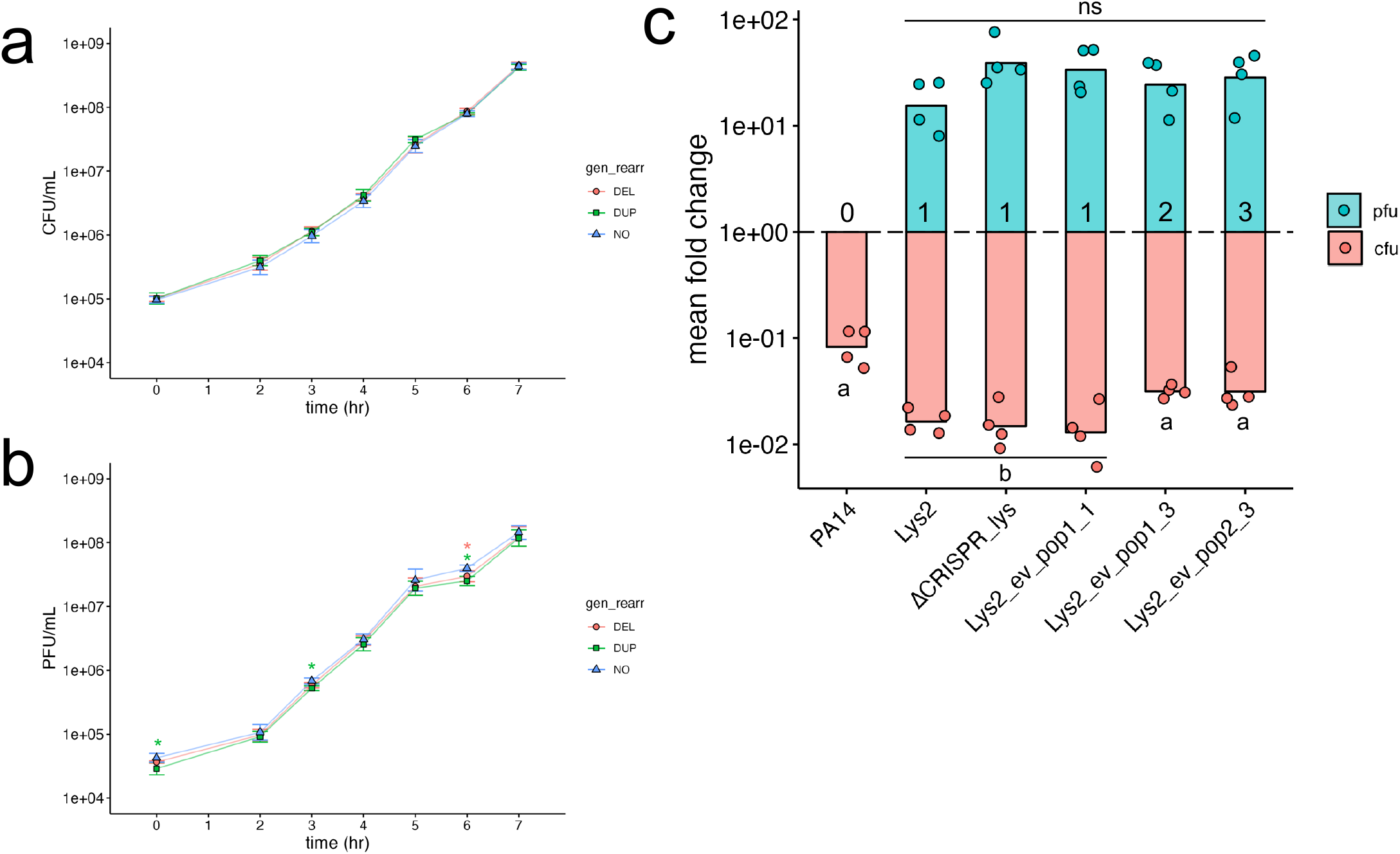
Genome rearrangements are tolerated without growth defects in rich medium in evolved lysogens. A) CFUs and B) PFUs of evolved lysogens in exponential growth. Time is indicated on the x-axis in hours, and CFUs or PFUs are indicated on the y-axis. Asterisks indicate significant differences by Dunn’s test between the genomic rearrangement (“DEL”: deletion; “DUP”: duplication) indicated by the color, and the “NO” genomic rearrangement group (the average of all other lysogens). C) Induction of lysogens with increasing numbers of viral genomes. Numbers in the PFU column represent how many viral genomes are present in the lysogen. Dunn’s Test of Multiple Comparisons with a Bonferroni correction was used to test the interaction between the mean fold change and the number of phages present. Points represent the mean of one experimental replicate (three technical replicates each). Bars represent the mean of all experimental replicates. Letters indicate statistical significance between groups; groups with the same letter are not statistically significant. “NS” = not significant.

**Figure S9.**
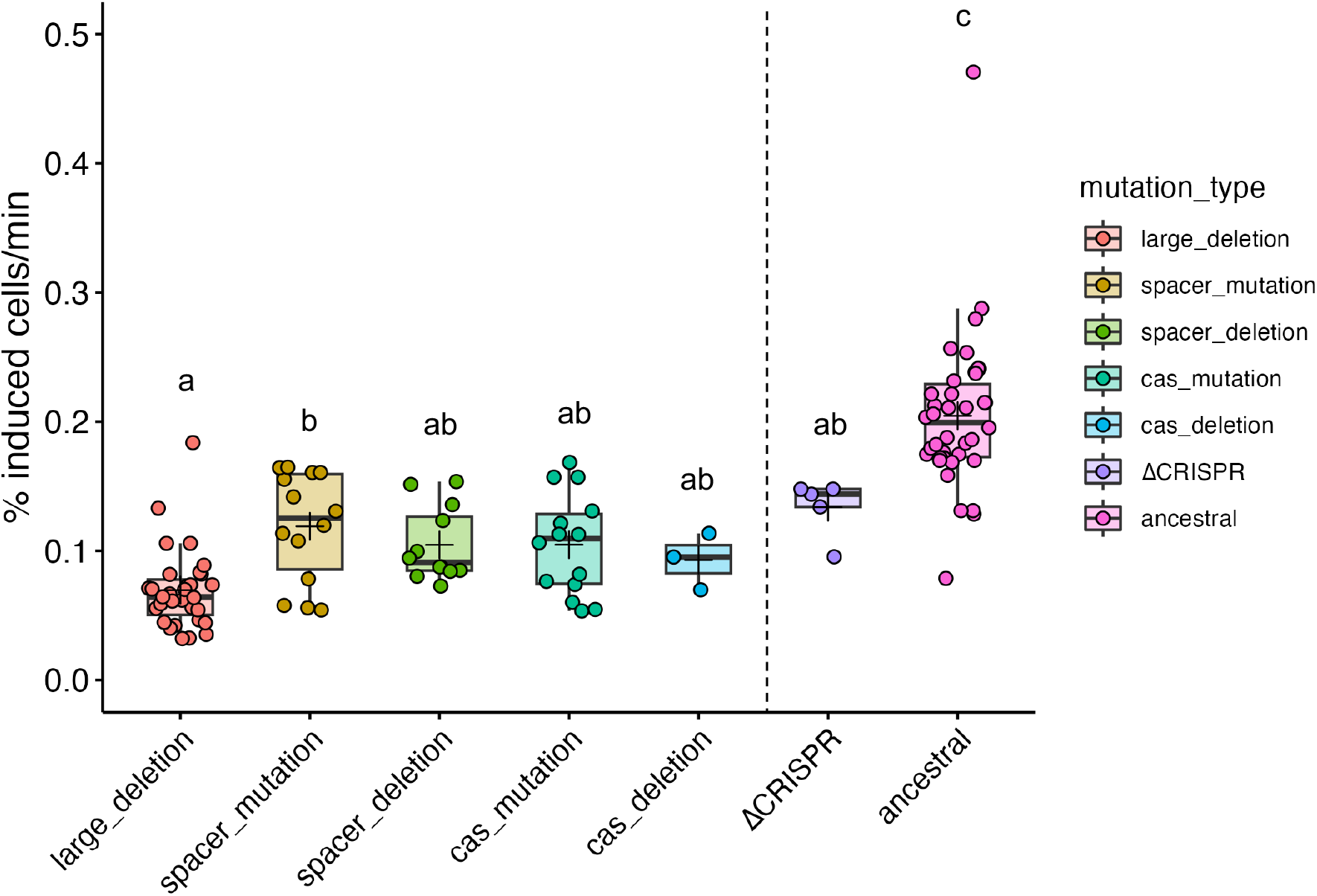
Spontaneous induction differences are maintained between groups and ΔCRISPR lysogen. Spontaneous induction was measured as above, except the growth curve was begun with a 1:100 diluton instead of 1:1000 (OD600 = 0.002). All differences between mutation groups was recovered, except the cas deletion group became significantly different from the ancestral, and the large deletion polylysogen group became significantly different from the spacer mutation group. Significance was tested with an ANOVA (*F_6_*_,110_ = 28.35, *P* < 2.2e-16) with a Tukey adjustment. Letters indicate significance; groups with different letters have a p-value <0.05; groups with the same or overlapping letters have a p-value of > 0.05. Points are the means of one experimental replicate from three technical replicates. A small jitter was added to the horizontal to increase visibility. Bars in the boxplots represent the median; crosses represent the means. Upper and lower bounds of the box are the upper and lower interquartile ranges.

**Table S1.** List of strains used in this study.

| Strain | Description | Source |
| --- | --- | --- |
| PA14 | WT *P. aeruginosa* strain | George O'Toole |
| Lys2 | PA14(DMS3) lysogen | Zegans et al, 2009 |
| PA14ΔCRISPR | CRISPR deletion | Cady et al, 2011 |
| PA14ΔCRISPR(DMS3) | CRISPR deletion lysogen | Cady et al, 2011 |

**Table S2.** Genes contained in Lys2 deletion. All functional predictions were confirmed by BLASTX using the protein RefSeq database.

| Lys2 Deletion Boundaries: 806165-826108 on PA14 Reference Genome |  |  |
| --- | --- | --- |
| Gene no. (5'-3') | Deleted gene | Description |
| 1 | PhzG | phenazine biosynthesis FMN-dependent oxidase |
| 2 | PhzF | phenazine biosynthesis protein; trans-2,3-dihydro-3-hydroxyanthranilate isomerase |
| 3 | PhzE | phenazine biosynthesis protein |
| 4 | PhzD | phenazine biosynthesis protein |
| 5 | PhzC | phenazine biosynthesis protein; phospho-2-dehydro-3-deoxyheptonate aldolase |
| 6 | PhzB | phenazine biosynthesis protein |
| 7 | PhzA | phenazine biosynthesis protein; |
| 8 | PhzM | phenazine-1-carboxylate N-methyltransferase |
| 9 | OpmD | multidrug efflux transporter outer membrane subunit; TolC family protein |
| 10 | MexI | MexW/MexI family multidrug efflux RND transporter permease subunit |
| 11 | MexH | MexH family multidrug efflux RND transporter periplasmic adaptor subunit |
| 12 | MexG | DoxX family protein; multidrug efflux RND transporter inhibitory subunit |
| 13 | PpgL | Gluconolactonase; 3-carboxymuconate cyclase |
| 14 | NmoR | LysR family transcriptional regulator |
| 15 | NmoA | nitronate monooxygenase |
| 16 | - | D-alanine--D-alanine ligase |
| 17 | - | MBL fold metallo-hydrolase |

**Table S3.** Description of mutations recovered in evolved uninfected PA14.

| Group | Sample ID | Mutated region | Description | Location on PA14-REF chromosome |
| --- | --- | --- | --- | --- |
| ev_pop1 | ev_pop1_1 | *fliF* | Nonsense mutation in flagellar M-ring protein G→A | 4459793 |
| ev_pop1 | ev_pop1_2 | *fliP* | Nonsynonymous mutation in start codon in flagellar biosynthesis protein C→T | 4072428 |
| ev_pop1 | ev_pop1_3 | *fliF* | Nonsense mutation in flagellar M-ring protein G→A | 4459793 |
| ev_pop1 | ev_pop1_4 | *fliF* | Nonsense mutation in flagellar M-ring protein G→A | 4459793 |
| ev_pop1 | ev_pop1_5 | *lasR* | Nonsynonymous mutation in transcriptional activator C→T | 4088064 |
| ev_pop1 | ev_pop1_6 | *fliF* | Nonsense mutation in flagellar M-ring protein G→A | 4459793 |
| ev_pop1 | ev_pop2_1 | *pilY1* | Frameshift mutation in T4P biogenesis factor protein GG→G | 5376474 |
| ev_pop1 | ev_pop2_2 | *pilY1* | Frameshift mutation in T4P biogenesis factor protein GG→G | 5376474 |
| ev_pop1 | ev_pop2_3 | *lasR* | Nonsynonymous mutation in transcriptional activator C→T | 4088064 |
| ev_pop2 | ev_pop2_3 | *pilY1* | Nonsynonymous mutation in T4P biogenesis factor protein T→A | 5378668 |
| ev_pop2 | ev_pop2_4 | *lasR* | Nonsynonymous mutation in transcriptional activator G→A | 4088043 |
| ev_pop2 | ev_pop2_4 | *pilY1* | Frameshift mutation in T4P biogenesis factor protein GG→G | 5376588 |
| ev_pop2 | ev_pop2_5 | *lasR* | Nonsynonymous mutation in transcriptional activator C→T | 4088064 |
| ev_pop2 | ev_pop2_5 | *pilY1* | 1kb deletion in T4P biogenesis factor protein | 5377966 |
| ev_pop2 | ev_pop2_6 | *lasR* | Nonsynonymous mutation in transcriptional activator G→A | 4088043 |
| ev_pop2 | ev_pop2_6 | *pilY1* | Frameshift mutation in type 4 pilus biogenesis factor protein GG→G | 5376588 |
| ev_pop2 | ev_pop3_1 | *fliF* | Nonsense mutation in flagellar M-ring protein G→A | 4459793 |
| ev_pop2 | ev_pop3_2 | *fliP* | Nonsynonymous mutation in start codon in flagellar biosynthesis protein C→T | 4072428 |
| ev_pop3 | ev_pop3_2 | MFS transporter | Synonymous mutation G→A | 4170934 |
| ev_pop3 | ev_pop3_3 | *fliF* | Nonsense mutation in flagellar M-ring protein | 4459793 |
| ev_pop3 | ev_pop3_4 | No mutations | NA | NA |
| ev_pop3 | ev_pop3_5 | C4 RNA | Point mutation in predicted C4 RNA | 4288779 |
| ev_pop3 | ev_pop3_6 | C4 RNA | Point mutation in predicted C4 RNA | 4288779 |

